# *pop-1/TCF*, *ref-2/ZIC* and T-box factors regulate the development of anterior cells in the *C. elegans* embryo

**DOI:** 10.1101/2021.12.14.472681

**Authors:** Jonathan D. Rumley, Elicia A. Preston, Dylan Cook, Felicia L. Peng, Amanda L. Zacharias, Lucy Wu, Ilona Jileaeva, John Isaac Murray

**Affiliations:** Department of Genetics, Perelman School of Medicine, University of Pennsylvania, Philadelphia, PA 19104; Division of Developmental Biology, Cincinnati Children’s Hospital Medical Center, Cincinnati, OH 45229; Department of Pediatrics, University of Cincinnati College of Medicine, Cincinnati, OH 45267

## Abstract

Patterning of the anterior-posterior axis is fundamental to animal development. The Wnt pathway plays a major role in this process by activating the expression of posterior genes in animals from worms to humans. This observation raises the question of whether the Wnt pathway or other regulators control the expression of the many anterior-expressed genes. We found that the expression of five anterior-specific genes in *Caenorhabditis elegans* embryos depends on the Wnt pathway effectors *pop-1/TCF* and *sys-1/*β-catenin. We focused further on one of these anterior genes, *ref-2/ZIC,* a conserved transcription factor expressed in multiple anterior lineages. Live imaging of *ref-2* mutant embryos identified defects in cell division timing and position in anterior lineages. *Cis-*regulatory dissection identified three *ref-2* transcriptional enhancers, one of which is necessary and sufficient for anterior-specific expression. This enhancer is activated by the T-box transcription factors TBX-37 and TBX-38, and surprisingly, concatemerized TBX-37/38 binding sites are sufficient to drive anterior-biased expression alone, despite the broad expression of TBX-37 and TBX-38. Taken together, our results highlight the diverse mechanisms used to regulate anterior expression patterns in the embryo.

**Graphical Abstract:** 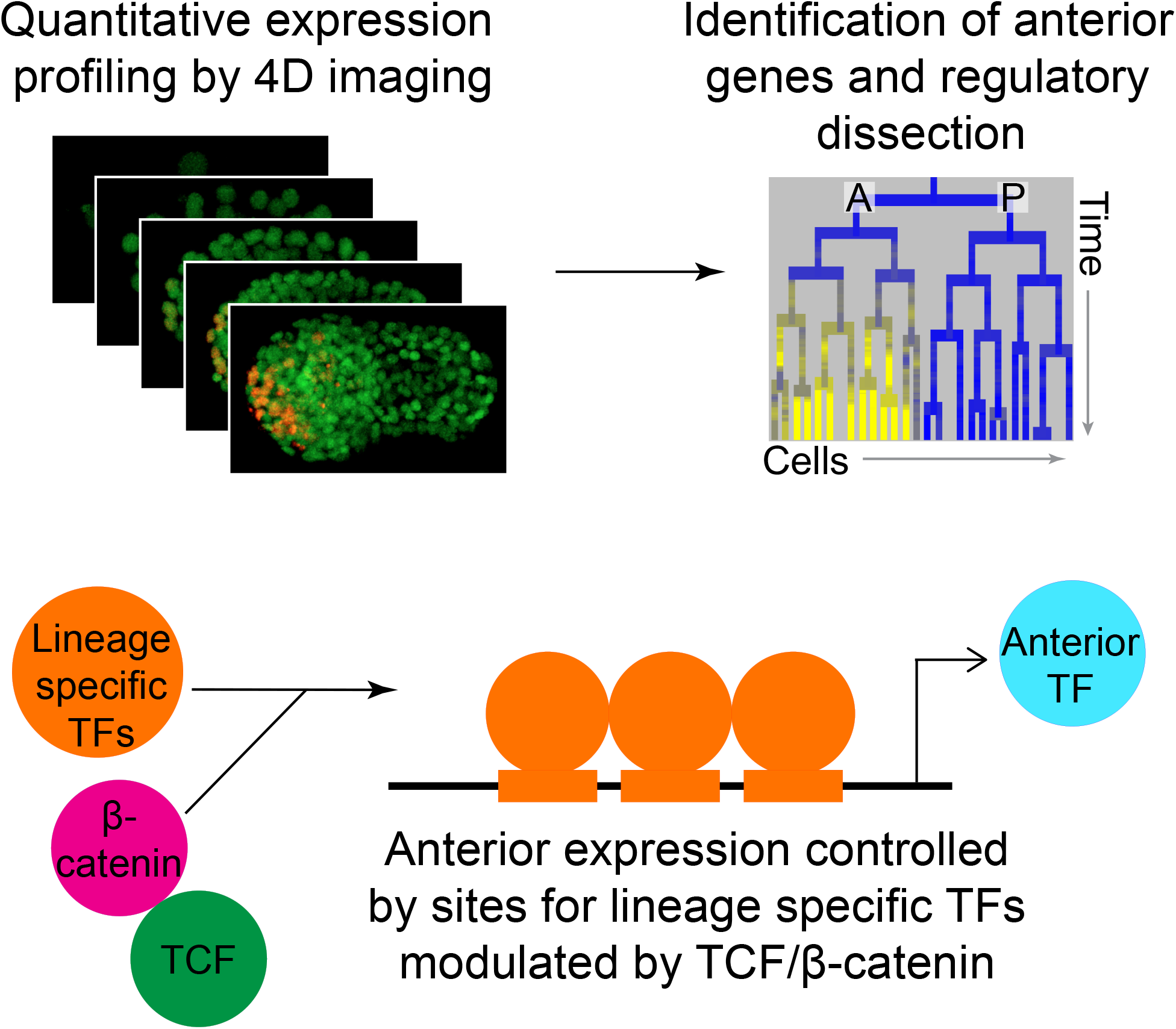

## INTRODUCTION

### Anterior-posterior patterning regulation

Proper anterior-posterior patterning is critical for animal embryonic development and requires the Wnt pathway across bilaterian animals from worms to humans. Typically, posterior cell identities are induced by posteriorly-produced Wnt ligands signaling through the canonical Wnt pathway [1]. In this pathway, signal transduction in response to Wnt activates posterior-expressed gene transcription through the transcription factor TCF and its co-activator β-catenin [2]. Although this conserved role for the Wnt pathway in regulating the expression of posterior genes is well documented [3], much less is known about how genes expressed in anterior cells are regulated.

*Caenorhabditis elegans* has been widely used to study anterior-posterior patterning because it is genetically tractable, its embryonic lineage is invariant, and most of its developmental regulators are highly conserved in vertebrates and other animals [4–10]. Also, the majority of embryonic cell divisions are oriented along the anterior-posterior axis, and require the Wnt pathway to differentiate the two daughter fates [11–13]. Following each anterior-posterior cell division, the Wnt pathway is activated in the more posterior daughter cell, leading to high nuclear SYS-1/β- catenin and low nuclear POP-1/TCF, whereas the anterior daughter cell has low nuclear SYS-1 and high nuclear POP-1[11–13]. The combination of high nuclear SYS-1 and low nuclear POP-1 in posterior cells makes it stoichiometrically favorable for POP-1 to be bound to posterior-expressed targets as a complex with SYS-1, thereby driving their expression [14–16]. In contrast, anterior cells have low nuclear SYS-1 and high nuclear POP-1; for posterior-expressed targets, this results in POP-1 being bound in the absence of SYS-1, allowing it to recruit co-repressors, including UNC-37/groucho [17] (Figure 1A). Loss of POP-1/TCF results in posterior genes having either reduced expression in posterior cells or increased expression in anterior cells, indicative of its dual roles as an activator and repressor. Loss of SYS-1/β-catenin results only in down-regulation of posterior genes in posterior cells, consistent with an exclusive role as a posterior activator [13]. These observations leave open the question of how anterior genes are regulated in *C. elegans*.

**Figure 1:**
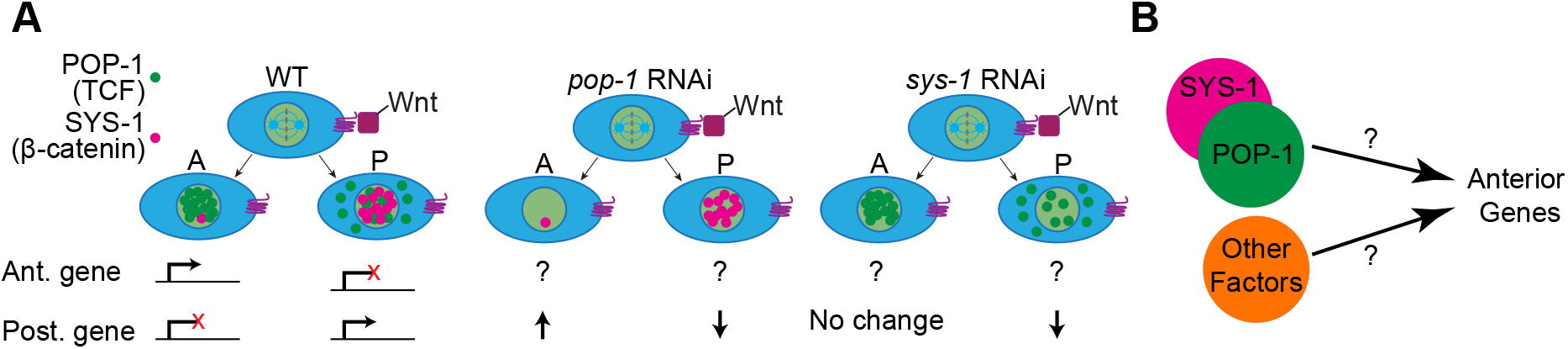
How are anterior genes regulated? A) The Wnt/β-catenin asymmetry pathway regulates anterior-posterior patterning by producing asymmetric nuclear concentrations of POP-1/TCF and SYS-1/β-catenin in anterior and posterior sister cells. Low nuclear concentrations of POP-1 and high nuclear concentrations of SYS-1 in the posterior cell activate expression of posterior genes and are associated with the lack of expression of anterior genes. Conversely, high nuclear concentrations of POP-1 and low nuclear concentrations of SYS-1 in the anterior cell repress posterior genes and are associated with expression of anterior genes. Effects of *pop-1* and *sys-1* RNAi on posterior genes are shown; effects on most anterior genes are unknown. B) In this work, we test whether *pop-1*/TCF and *sys-1*/β-catenin regulate anterior genes in the *C. elegans* embryo, and ask what other factors also regulate anterior expression.

Previous work suggests at least three possible mechanisms by which anterior genes could be regulated. 1) They may be regulated by the Wnt pathway indirectly, with posteriorly-expressed canonical Wnt targets repressing the expression of anterior genes in posterior cells. 2) They may be directly regulated by Wnt pathway components acting in a modified (or “opposite”) manner. 3) They may be regulated by a non-Wnt-related mechanism (Figure 1B).

An example of indirect Wnt pathway regulation through a posterior repressor is seen in the *C. elegans* embryonic EMS lineage, in which cells derived from the posterior daughter of EMS, E, express the POP-1-activated, gut-specifying transcription factor *end-1*. In turn, *end-1* represses the transcription factor *ceh-51*, which is normally expressed in and specifies the lineage derived from the anterior daughter of EMS, MS [18,19].

Conversely, the *C. elegans* anterior gene *ttx-3* appears to be directly regulated by Wnt pathway components. *ttx-3* helps specify the AIY neuron class and is regulated by POP-1 and SYS-1 in a manner dependent on the sole *C. elegans* ZIC family transcription factor REF-2 [20]. After the AIY grandmother divides, *ttx-3* is expressed in the anterior daughter, the AIY mother, but not in the posterior daughter. In the AIY mother, the Wnt pathway is inactive, with low nuclear SYS-1 and high nuclear POP-1, and both *pop-1* and *ref-2* are required to activate *ttx-3* expression. In the posterior sister, the Wnt pathway is active, with high nuclear SYS-1 levels and low nuclear POP-1 levels, and *sys-1* is required to repress expression of *ttx-3*. POP-1 and REF-2 can directly interact, suggesting that REF-2 and POP-1 bind as a complex to activate *ttx-3* expression in the AIY mother. The role of SYS-1 is less clear but some evidence suggests SYS-1 may bind to the POP-1/REF-2 complex to repress expression, or may sequester away the limited POP-1, such that POP-1 cannot interact with REF-2 [20,21]. Similar regulation has also been seen in other species. For example, *Drosophila* TCF/Pangolin can act in a “reverse transcriptional switch” such that when TCF is bound to a non-optimal site without binding to β-catenin/Armadillo it activates expression, and when it binds along with β-catenin it represses expression [22].

Because of the role of POP-1 and SYS-1 in regulating the anterior-specific expression of *ttx-3*, we hypothesized that these Wnt pathway components may also regulate other anterior genes. POP-1 and SYS-1 could act as general anterior expression regulators, interacting with lineage-restricted co-regulators to ensure appropriate expression of anterior genes in the correct lineages [20,21]. We tested this by mining single-cell-resolution expression data to identify transcription factors expressed in anterior-specific patterns [23–25], and show that of five we tested, all require *pop-1* and/or *sys-1* for either anterior expression or posterior repression. We focus in more detail on a single anterior-specific gene, *ref-2/ZIC*. By automated lineage analysis of mutants, we find that *ref-2* is required for normal cell division timing and cell position in *ref-2* expressing lineages. Embryonic expression of *ref-2* is driven by at least three developmental enhancers, two of which drive early embryonic expression and one of which drives expression in later-stage embryos. The most distal enhancer (3.9 kb upstream of the transcription start site) drives highly anterior-biased expression. Functional dissection revealed striking redundancy within this enhancer, reminiscent of enhancer redundancy that facilitates robust gene expression in other species [26]. We also identified a role for the T-box transcription factors *tbx-37* and *tbx-38* in driving anterior-specific expression. Surprisingly, concatemers of a single T-box binding site reiterate much of the anterior-specific expression pattern of *ref-2* in early embryos, suggesting a key role for these genes in driving anterior expression.

## RESULTS

### Many early embryonic transcription factors have anterior-specific expression

We used large-scale single-cell expression databases derived from time-lapse 4D imaging of embryos expressing reporter genes to identify anterior-expressed genes (Figure 2A). We define “anterior” or “posterior” lineages as sets of related cells derived from either the more-anterior or more-posterior daughter after a specific cell division (Figure 2B). Notably, this is distinct from physical position; for example, a cell that divides near the posterior pole of the embryo gives rise to both an anterior and posterior daughter lineage, both of which are located in the posterior half of the embryo (Figure 2B). Following convention [5,27], all lineage trees in this manuscript are drawn with the more anterior daughters on the left and the posterior daughters on the right of each bifurcation. Earlier studies identified a strong tendency for genes to be expressed in either multiple anterior or multiple posterior lineages during early and mid-embryogenesis; these frequently appear as “lineally repetitive” patterns, with multiple related anterior or posterior cousin lineages all expressing the same gene (e.g. *nob-1* and *ref-2* in Figure 2D, E) [23,24]. Both anterior-specific and posterior-specific gene expression were similarly common. While posterior specific genes (e.g. Figure 2D) appear to be largely regulated by canonical Wnt signaling [13], it is less clear how anterior-specific patterns are regulated. We mined existing literature and large-scale databases of fluorescent reporter expression patterns (Figure 2A) [23–25] to identify 19 conserved genes expressed in anterior lineage-specific patterns (Figure 2C).

**Figure 2:**
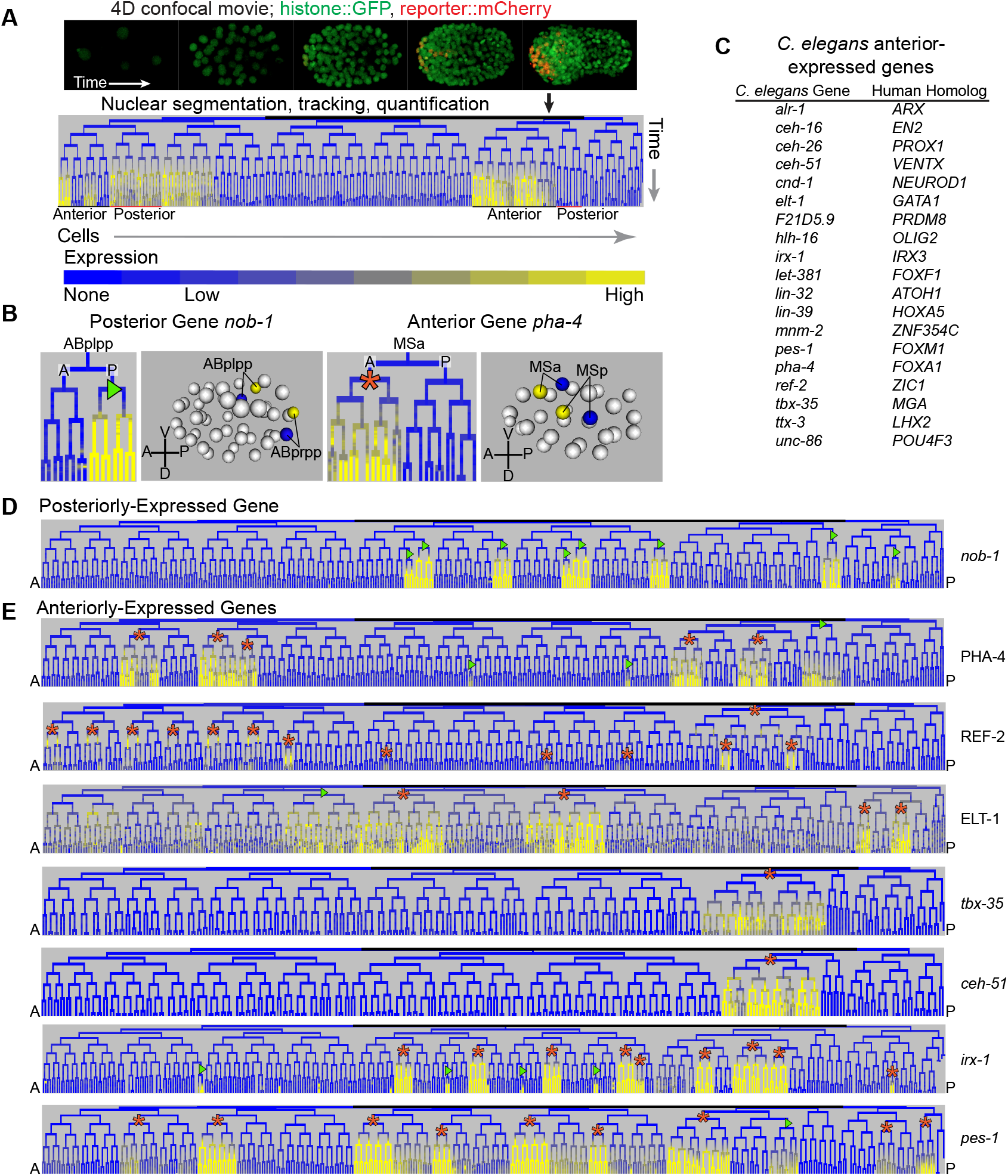
Identification of genes with anterior-biased expression. A) Automated lineaging traces lineages and quantifies expression from 4D confocal images. In the lineage trees, vertical lines represent cells, and horizontal lines represent cell divisions. Most cells divide along the anterior-posterior axis, with anterior cells depicted on the left branch and posterior cells on the right branch. B) Posterior and anterior genes are denoted as such based on their expression in cells descended from either a posterior or anterior sister cell, respectively, following an anterior-posterior cell division. Anterior and posterior genes can generally be expressed in cells from any part of the embryo. Anterior and posterior founder cells are labeled with orange asterisks/green triangles respectively. C) List of anterior-biased genes in the EPiC database along with their predicted human homologs. D) Expression pattern of a posterior gene, *nob-1/Hox9-13*. E) Expression patterns of seven anterior genes; most are expressed in unique combinations of mostly anterior lineages.

We measured cellular resolution expression profiles of seven of these by live confocal imaging followed by automated cell tracking and lineage tracing (Figure 2A, E, Figure S1, Figure S2, Figure S3) [25,28]. For each gene, we collected 4D time-lapse images of transgenic embryos expressing a histone-mCherry (*pha-4*, *ref-2*, *elt-1*, *tbx-35*, *irx-1*, and *pes-1*), or GFP (*ceh-51*) reporter under the control of upstream regulatory sequences of each gene (referred to here as “promoter reporters”). For five genes we also collected images for fosmid transgenes (*irx-1, elt-1, pha-4* and *ref-2)* or CRISPR knock-ins (*ceh-51*) in which the protein is fused to GFP and surrounded by its normal genomic context (“protein reporters”). Each embryo also contained a ubiquitously expressed second-color, histone-GFP or histone-mCherry fusion, used for cell tracking. We identified each nucleus at each time point and traced them across time using StarryNite [28,29] cell tracking software, and used AceTree [30,31] for subsequent manual error correction and validation that the extracted lineages were correct. Finally, we quantified the expression of the reporter in each nucleus at each time point (Figure 2A) [25]. The results confirm the anterior-specific expression of each gene, and, in some cases, identify previously unknown expression patterns (Figure S1, Figure S2, Figure S3). For each promoter reporter, the anterior-specific patterns were consistent, but the protein fusion reporters all had additional expressing lineages and dynamics indicative of distal enhancers and post-transcriptional regulation (Figure S1, Figure S2, Figure S3). The anterior-biased expression of these genes is also seen in embryonic single-cell RNA-seq data (Figures S4 and S5) [32].

### The Wnt pathway effectors *pop-1* and *sys-1* are required for normal anterior-specific expression

Most genes expressed preferentially in posterior lineages require the Wnt effector transcription factor POP-1/TCF either for activation in posterior cells (together with its coactivator SYS-1/β-catenin) or for repression in anterior cells, and at least some are direct targets [12,13,33–36]. Although *pop-1* is required for the normal expression of some anterior genes [20,33,37], it is unclear whether *pop-1* and *sys-1* are generally required for anterior-specific expression, as they are for posterior-specific expression.

To test this, we measured the expression of five anterior gene reporters (*tbx-35*, *ceh-51*, *elt-1*, *irx-1*, and *ref-2*) after *pop-1* and *sys-1* RNAi by live 4D imaging and lineage tracing as described above. We used promoter reporters for *tbx-35*, *ceh-51*, *irx-1*, and *ref-2* because our wild-type expression data indicated these reporters are sufficient for anterior regulation (Figure S1, Figure S2, Figure S3). For *elt-1*, anterior-specific expression was more robust for the protein reporter so we tested that reporter’s dependence on *pop-1* and *sys-1* (Figure 3; Figure S6).

**Figure 3:**
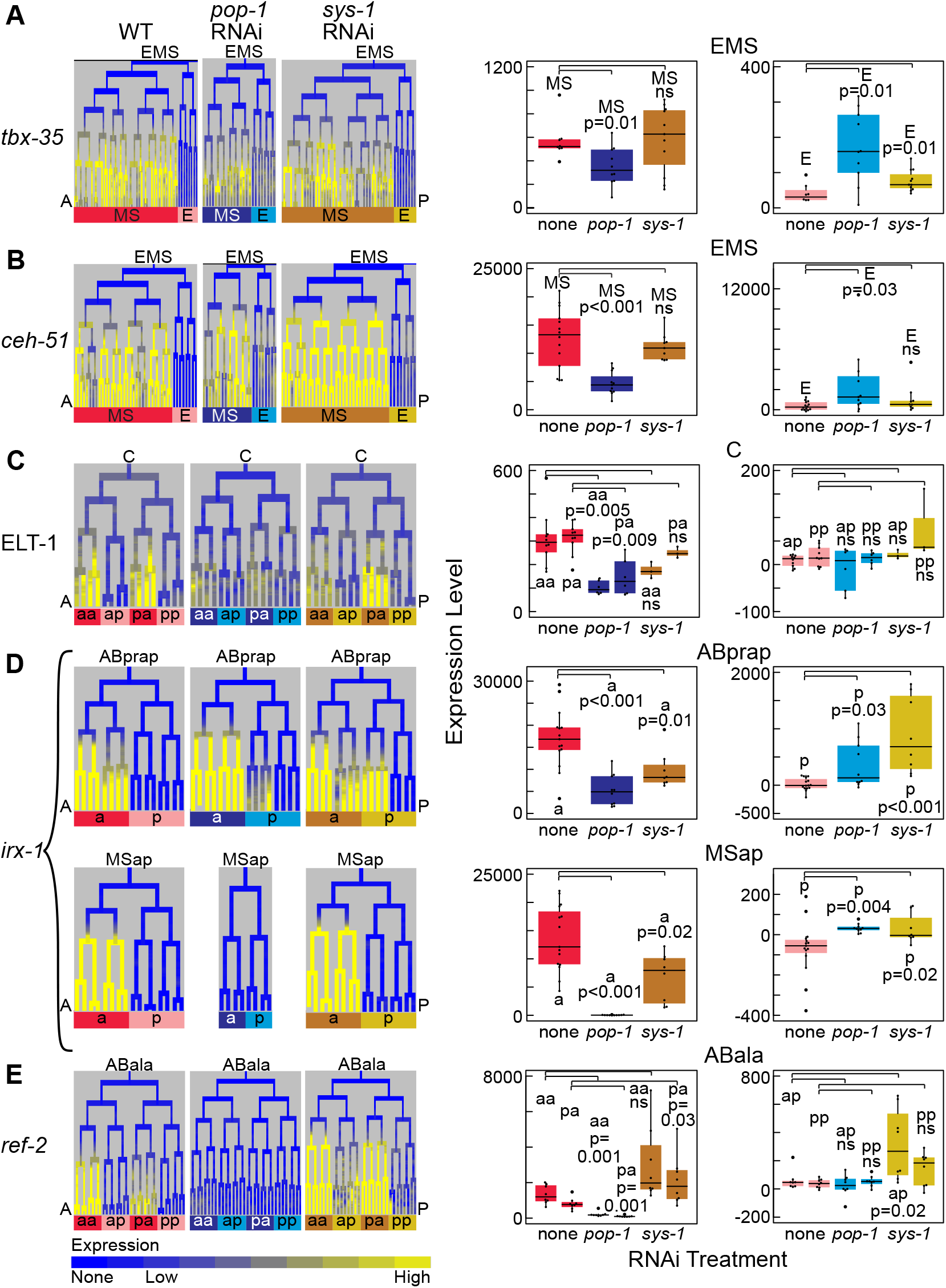
Regulation of anterior genes by Wnt effectors *pop-1* and *sys-1*. A-E) Expression pattern (left) and quantification (right) of reporters for the anterior genes *tbx-35* (A), *ceh-51* (B), ELT-1 (C), *irx-1* (D), and *ref-2* (E) in specific anterior lineages and their posterior sisters. Expression is shown under WT conditions and following *pop-1* or *sys-1* RNAi. Expression quantification (right) is the mean expression across all measurements within that lineage from when the reporter is normally first detected until the last time point indicated in the lineage (details in Methods).

Reporters of two of the genes we tested, *tbx-35* and *ceh-51*, are expressed solely in the anterior lineage MS (Figure S2; Figure S6A, B), and previous qualitative expression analyses indicated that expression persists in the absence of *pop-1* [19,37,38]. Consistent with this, we detected expression of both genes in the MS lineage after *pop-1* RNAi, however this expression was significantly decreased. We also observed increased expression of both genes in the E lineage (posterior sister of MS), although this expression remained lower than both the wild-type and *pop-1* RNAi MS levels (Figure 3A, B; Figure S6A, B; Table S4). We also observe low-level ectopic expression of *tbx-35* in the E lineage under *sys-1* RNAi. These results indicate that both *tbx-35* and *ceh-51* require *pop-1* for full expression in the MS lineage and for repression in the E lineage (as shown previously [19,37]), and that *sys-1* is required to repress *tbx-35* in the E lineage.

*elt-1* reporters are expressed in several early anterior lineages that primarily give rise to ectodermal fates (ABpla, ABpra, Caa and Cpa), and in one posterior lineage (ABarp) (Figure S1; Figure S6C). In the Caa and Cpa lineages, anterior lineages that express both the promoter and protein reporters, *pop-1* RNAi results in decreased *elt-1* reporter expression. However, RNAi for neither *pop-1* nor *sys-1* affects expression in their posterior sister lineages Cap and Cpp (Figure 3C; Figure S6C; Table S4). Similarly, *pop-1* RNAi results in reduced expression of the protein reporter in the ABpla and ABpra lineages, while RNAi knockdown of neither *pop-1* nor *sys-1* has much effect on their posterior sister lineages ABplp and ABprp (Figure S6C; Table S4). Thus, in both the ABp and C lineages, *pop-1* is required for anterior expression of *elt-1*, but neither *pop-1* nor *sys-1* is required for posterior repression.

*irx-1* reporters are expressed early in four related anterior sublineages of ABp, and in three anterior sublineages of MS, while in later embryos they are expressed in both anterior and posterior lineages (Figure S3; Figure S6D). Like for *elt-1, pop-1* RNAi reduces *irx-1* reporter expression in anterior ABp-derived sublineages, and two of the MS-derived expressing sublineages lose expression nearly completely. After both *pop-1* and *sys-1* RNAi, the *irx-1* reporter expression expands into several posterior sublineages of ABp whose anterior sisters normally express *irx-1* (Figure 3D; Figure S6D; Table S4). These results indicate that *irx-1* requires *pop-1* for expression in anterior lineages and both *pop-1* and *sys-1* for repression in posterior lineages.

Reporters for *ref-2* are expressed in the early embryo in all MS descendants and in several related anterior sublineages of ABa that arise at the 50-cell stage (Figure S1; Figure S6E). In later embryos, expression occurs in two anterior sublineages of MS, in several anterior ABp-derived sublineages, and in the Pn ventral epidermal blast cells, consistent with previous studies [39]. After *pop-1* RNAi, expression of the *ref-2* promoter reporter is significantly reduced in anterior sublineages of ABala (Figure 3E). As for other genes, *sys-1* RNAi caused de-repression of *ref-2* in several posterior sisters of the normally expressing sublineages, but we did not detect de-repression in these sublineages after *pop-1* RNAi (Figure 3E; Figure S6E; Table S4). Thus, *ref-2* requires *pop-1* for expression in anterior lineages and *sys-1* for repression in posterior lineages.

To summarize these perturbations, each anterior gene tested requires *pop-1* for full expression in anterior lineages. Several genes also require *pop-1* (*tbx-35*, *ceh-51* and *irx-1*) or *sys-1* (*tbx-35, irx-1* and *ref-2*) for repression in some posterior lineages. Thus, it appears that the expression of most anterior genes requires the Wnt pathway components POP-1 and SYS-1, but this dependency is complex as was seen previously for posterior genes [13]. Another layer of complexity is that in some cases, we cannot rule out that the loss of expression after *pop-1* RNAi is, at least in part, due to indirect effects of *pop-1* loss, such as an earlier anterior to posterior fate conversion. For example, for *irx-1* expression in descendants of the MS lineage, an MS to E fate conversion could explain the loss of expression after *pop-1* RNAi. However, in this case, we observe consistent reduction in *irx-1* reporter expression after *pop-1* RNAi even in embryos without a strong MS to E conversion (as assayed by cell cycle timing). Expression changes of *irx-1* in the AB lineage cannot be explained by known fate conversions, indicating that the expression changes are at least in part due to a direct effect of *pop-1*.

### The anterior-specific transcription factor *ref-2/ZIC* is required for normal patterns of cell division and cell position in expressing lineages

The striking anterior-specific expression patterns observed for early embryonic TFs raises the question of whether these factors regulate the development of anterior lineages. To address this question, we focused on one anterior gene, *ref-2*. *ref-2* encodes a zinc finger transcription factor homologous to mammalian ZIC genes and *Drosophila* Odd-Paired. Previous work showed that *ref-2* is required to prevent progeny of “Pn” epidermal blast cells from fusing with the major hypodermal syncytium in larval stages [39] and a predicted null allele of *ref-2*, *ref-2(gk178)*, causes larval lethality [40]. Recently, REF-2 was shown to activate the anterior-specific expression of the neural regulator *ttx-3* in the AIY mother cell by binding the *ttx-3 cis-*regulatory region and recruiting POP-1 independent of POP-1 DNA binding activity [20]. Since *ref-2* is expressed in many other anterior lineages in early embryos, we investigated whether *ref-2* mutants have developmental defects in those lineages.

We tested for embryonic lineage phenotypes at single-cell resolution by analyzing 4D time-lapse images of eight embryos homozygous for the deletion allele *ref-2(gk178)* and expressing a ubiquitous histone-mCherry fusion for cell tracking. We used StarryNite to track the lineage of each embryo, and used AceTree to curate the lineages through the >550-cell stage (six embryos) or >350-cell stage (two embryos). We compared the position and cell cycle length of each cell in each embryo to those from a reference of 17 wild-type embryos [41,42] to identify outliers (“defects”) (Figure 4A).

**Figure 4:**
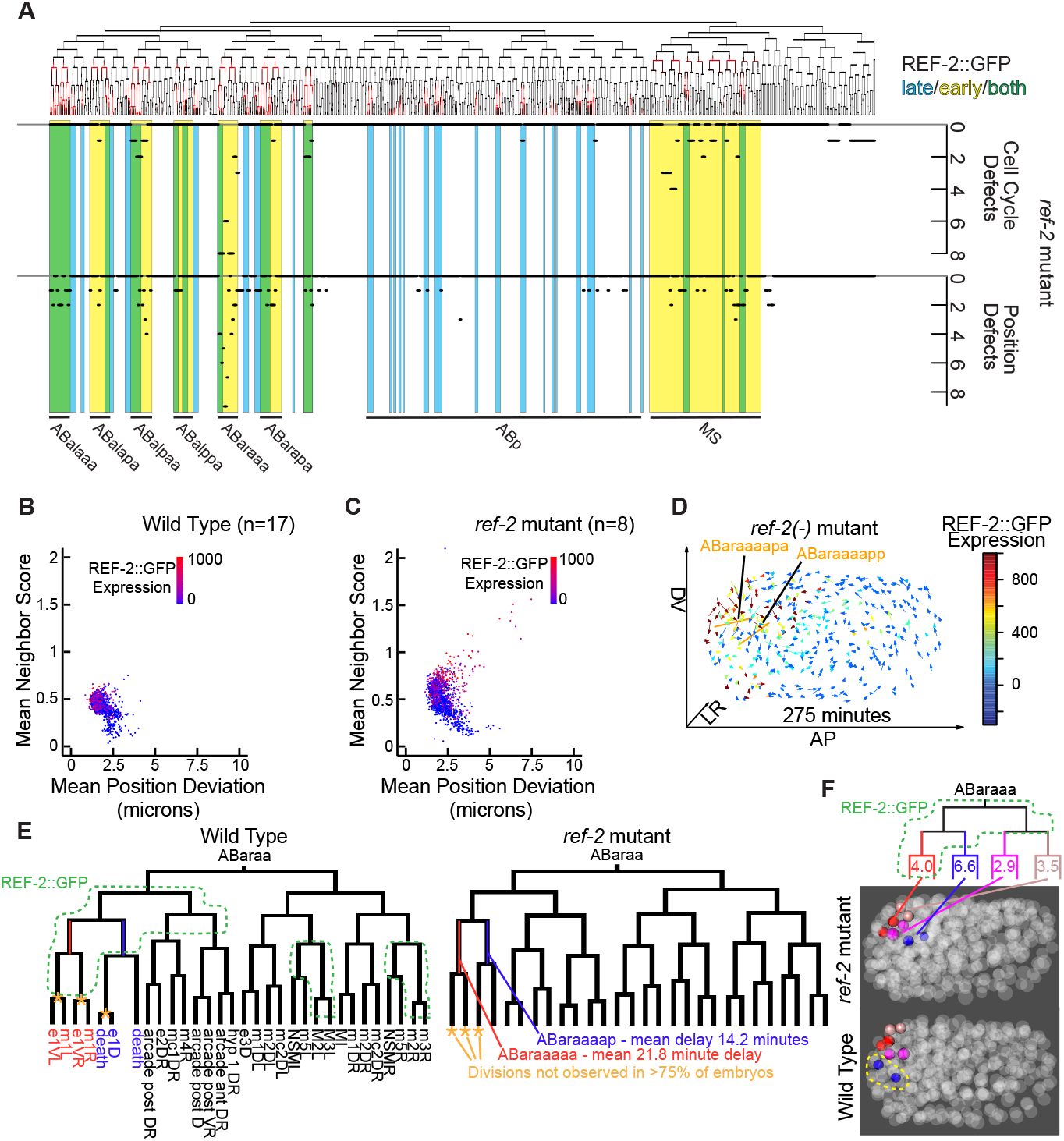
*ref-2* is required for proper division timing and positioning for many embryonic cells that are descended from cells that express *ref-2*. A) Lineage tree depicting the REF-2 protein expression pattern (top), and chart indicating the number of embryos out of eight homozygous *ref-2* mutant embryos that exhibit cell cycle defects and position defects in indicated cell lineages. Yellow indicates lineages that express *ref-2* early, blue indicates lineages that express *ref-2* late, and the overlap (green) indicates both. A corresponding analysis of wild-type embryos gives 0-1 defects per cell out of all 17 embryos tested [41]. B, C) Scatter plot of mean neighbor score vs mean position deviation of cells of 17 wild-type (B) and eight *ref-2* mutant (C) embryos. Neighbor score is the ratio of each cell’s distances to its ten closest wild-type neighbors between mutant and wild-type embryos. Points representing cells are colored based on WT REF-2::GFP expression levels [41]. D) Plot of cell position deviations with arrows starting at the average WT position of cells and pointing to the average position in *ref-2* mutants. Arrows are colored by the WT expression levels of REF-2::GFP for each cell. The labeled cells (ABaraaaapa and ABaraaaapp) have the greatest mean cell position deviations. E) ABaraa lineages of WT and *ref-2* mutant embryos, with cells expressing REF-2::GFP outlined on the WT tree. Two delayed and three missed cell divisions are highlighted on the *ref-2* mutant tree. F) ABaraaa lineage with cells expressing REF-2::GFP outlined. The average position deviations in microns of terminal sister cells are indicated on the terminal branches of the lineage tree. Three dimensional projections of a WT and a *ref-2* mutant embryo are shown with the positions of the terminal ABaraaa lineage cells highlighted. ABaraaaapa and ABaraaaapp are outlined in the WT embryo projection as the cells with the greatest mean position deviation in the *ref-2* mutant.

We identified a total of 55 cells with early, late or missing divisions, and 54 of these were derived from REF-2::GFP expressing lineages (P < 10^-15^). Overall 95% of cell cycle defects were delays or missed divisions, indicating a role for *ref-2* in promoting cell division. Similarly, we identified 68 position defects, of which 56 were in *ref-2* expressing lineages (Figure 4B, C, P = 1.1 * 10^-14^, chi-squared test). Ten *ref-2-*expressing cells, and no non-expressing cells, had position defects in two or more embryos. The cells with the strongest position and cell cycle length defects were from the *ref-2*-expressing anterior ABaraaa lineage, which produces primarily anterior pharyngeal cells (e1, m1 and arcade cells). The most frequent cell cycle length defects were in ABaraaaaa and ABaraaaap (both defective in 8 of 8 embryos), which were delayed by 21.8 minutes and 14.2 minutes, respectively (Figure 4E). The largest position defects were in ABaraaaapa and ABaraaaapp, which were mispositioned on average 6 microns posterior of their wild-type position (Figure 4D, F). However, other defects were broadly distributed across *ref-2-*expressing lineages, in particular MS- and ABa-derived anterior sublineages (Figure 4A). In summary, the anterior lineage-expressed TF *ref-2* is required for the normal development of anterior lineages, although the low penetrance of many defects suggests that *ref-2* may act with other partially redundant regulators, similar to other early zygotic TFs [41,43–48].

### Modular enhancers control *ref-2* embryonic expression

The anterior expression and phenotypes of *ref-2* raise the question of what sequences and regulators control this expression. To identify genomic sequences responsible for *ref-2* anterior expression, we first compared expression driven by the 31.9 kb genomic fosmid protein reporter to the shorter 4.1kb upstream (“promoter”) reporter (Figure 5A, B, C; Figure S8). In general, the patterns were similar; both are expressed broadly in the MS lineage and in multiple anterior ABa-derived sublineages. In the ABa lineage, the promoter reporter shows background expression in some posterior sister sublineages suggesting the existence of repressive sequences outside of this promoter region. Additionally, the promoter fusion reporter is detected persistently in both the MS and ABa lineages, whereas protein reporter expression is detected transiently, likely reflecting the use of a stable histone-mCherry reporter for the promoter reporter. Importantly, both the REF-2::GFP protein and *ref-2* promoter reporters drive expression in progenitors of most cells defective in *ref-2* mutants.

**Figure 5:**
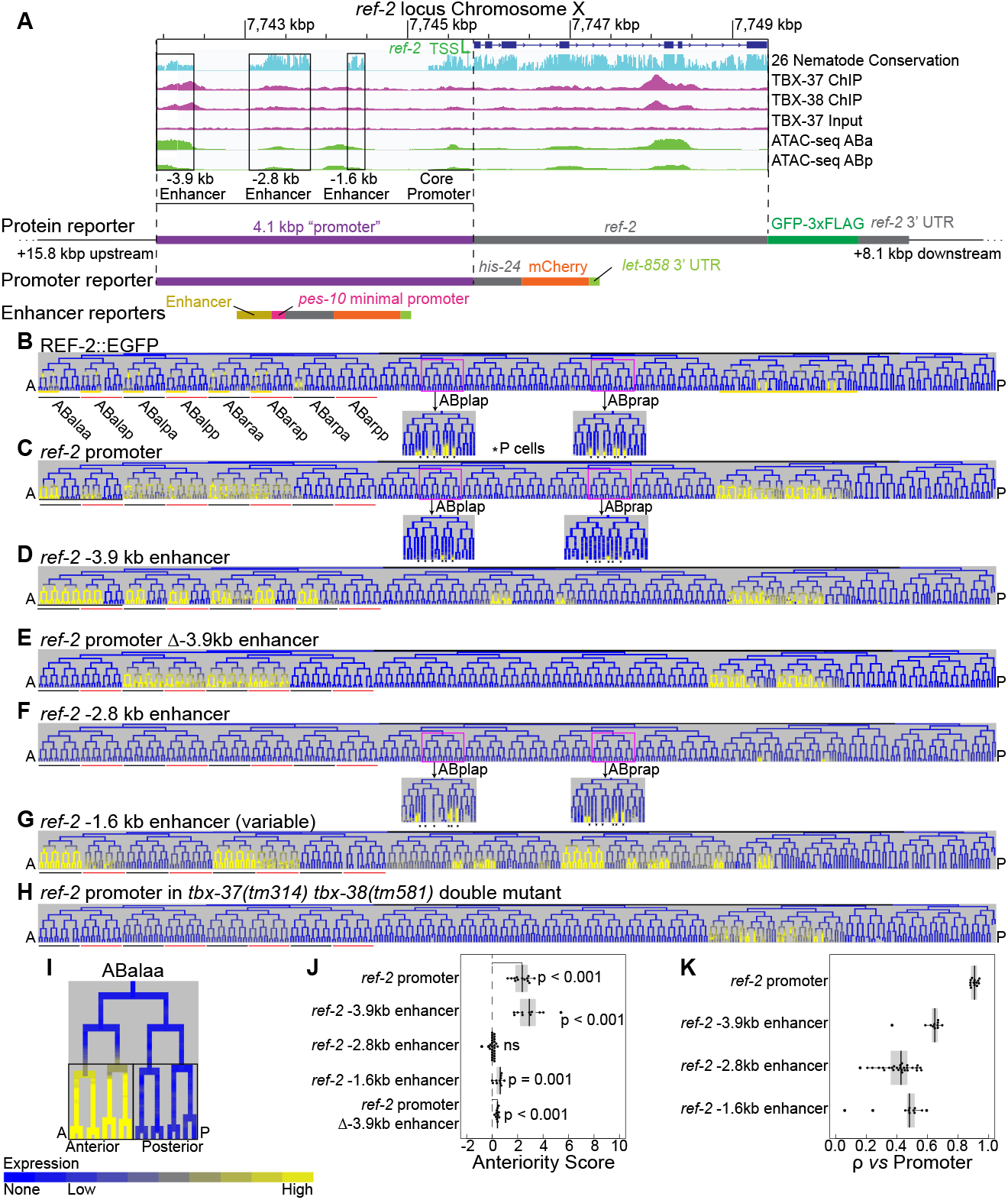
Identification of *ref-2* enhancers. A) Genome browser screenshot showing the *ref-2* locus on the X chromosome. Displayed tracks include the gene model, sequence conservation among 26 *Caenorhabditis* species [49], ChIP-seq data for TBX-37 and TBX-38 binding, and ATAC-seq data for cells in the ABa and ABp lineages [50]. Boxes indicate candidate enhancer regions as determined by conservation. Below the genome browser screenshot are models of the fluorescent reporter expression constructs we examined. B-H) Lineage trees showing the expression patterns of *ref-2* protein (B), promoter (C), −3.9 kb enhancer reporter (D), promoter lacking the −3.9 kb enhancer (E), −2.8 kb enhancer reporter (F), −1.6 kb enhancer reporter (G), and promoter in the *tbx-37 tbx-38* double mutant (H) as determined by time-lapse confocal microscopy. Sublineages used in anteriority analyses are labeled in B and underlined in black (anterior) and red (posterior) under each lineage tree. Sublineages with transient early REF-2::EGFP (protein reporter) expression are underlined in yellow under the lineage tree in B. Pn ventral epidermal cells are indicated with asterisks in insets of panels B, C, and F. I) Example lineage showing which cells were included in anteriority score analyses. Reporter expression levels were averaged across cells in the anterior or posterior lineages for the indicated cell generations. J) Anteriority scores of the *ref-2* promoter, enhancers, and promoter lacking the −3.9 kb enhancer. Lineages used for analysis are ABala, ABalaa, ABalap, ABalpa, ABalpp, ABaraa, ABarap, ABarpa, and ABarpp (see Figure 5B). K) Full somatic lineage correlation (Spearman’s ρ) of each reporter’s expression pattern with the mean expression pattern across embryos expressing the promoter reporter.

To identify the sequences in this region that regulate expression in these lineages, we sought to identify minimal portions of the region with enhancer activity. We identified three portions of the *ref-2* promoter conserved among *Caenorhabditis* species, suggesting they encode important functions (Figure 5A) [49,51,52]. The most distal of these regions is in open chromatin in ABa-derived cells as measured in a recently published ATAC-seq data set [50]. Also, this region is bound by the broadly-expressed ABa lineage transcription factors TBX-37 and TBX-38, as determined by a recently published ChIP-seq data set (Figure 5A) [50]. We fused each putative enhancer to a reporter cassette consisting of a minimal *pes-10* promoter and a *his-24*::mCherry reporter. The *pes-10* promoter is widely used, drives no consistent embryonic expression on its own [13,53,54] and is compatible with a wide variety of enhancers [13,53]. We introduced each reporter into worms and used StarryNite to determine in which cells each putative enhancer drives expression.

The most distal enhancer, located 3.9 kb upstream of the likely transcription start site, (−3.9 kb enhancer; 449 base pairs) drives early expression (during gastrulation) in several anterior sublineages of ABa and in anterior sublineages of MS (Figure 5D; Figure S8). This expression occurs in nearly identical lineages to the early portion of the protein reporter expression pattern, but while the protein reporter expression in these lineages is transient, the enhancer reporter persists, again likely because of the use of a stable histone-mCherry reporter. The −3.9 kb enhancer expression is also well correlated with the pattern driven by the full promoter of *ref-2* (Spearman’s ⍴ > 0.6, Figure 5K). Deleting this enhancer from the *ref-2* promoter reporter results in loss of the strong anterior-biased expression in the ABalaa, ABalap, and ABarpa lineages, and a loss in the weak anterior-biased expression in the ABalpa, ABalpp, ABaraa, and ABarap lineages. All remaining consistent expression occurs in the lineages derived from the Notch-signaled cells ABalp and ABara and has little anterior bias (Figure 5E, J; Figure; S7; Figure S8) [48]. The significance of this expression is unclear; it can be seen in the promoter reporter but not the protein reporter, suggesting that it is normally repressed by additional sequences outside the promoter. We conclude that the −3.9 kb enhancer is necessary and sufficient for anterior-biased expression in the ABa lineage.

A second 745 bp enhancer, located at −2.8 kb, drives expression in later embryos (bean stage) in some anterior sublineages of MS and in the ABp-derived Pn ventral epidermal blast cells that were previously reported to require *ref-2* [39]. These cells also robustly express the *ref-2* protein reporter and weakly express the *ref-2* promoter reporter (Figure 5B, C, F, J, K). Finally, the most proximal enhancer candidate (at −1.6 kb; 206 base pairs) drives variable, weakly anterior-biased, expression in multiple lineages, most consistently in ABala and ABara (Figure 5G, J, K; Figure S9) and also drives late expression in some ABp sublineages (Figure 5G; Figure S9). Much of this expression is in cells or at stages that do not express the full-length reporters, suggesting that the activity of this sequence differs in its normal genomic context.

To measure anterior expression bias, we developed an “anteriority score” based on the log2 ratio between anterior and posterior sister lineage expression. Because expression tends to accumulate over several cell cycles in each expressing lineage, we calculated this score based on the mean expression of the progeny 2-3 cell divisions after the birth of each anterior lineage (Figure 5I). Using this metric, we detected robust and significant anterior-biased expression for both the *ref-2* promoter and −3.9 kb enhancer reporters in ABa-derived sublineages, and reduced or no anterior-biased expression for the −2.8 kb and −1.6 kb enhancer reporters and for the promoter Δ −3.9 kb enhancer reporter in the same sublineages (Figure 5J). Comparing the expression driven by each enhancer to the full promoter showed each is positively, but imperfectly, correlated, consistent with each driving a subset of the full pattern (Figure 5K).

### *ref-2* promoter expression requires the T-box transcription factors *tbx-37* and *tbx-38*

A dominant feature of the early *ref-2* expression pattern is reiterated expression in six of the eight anterior sister cells of the ectodermal ABa lineage at the 50-cell stage (AB32 stage--when there are 16 ABa descendants). This expression is first observed in the fluorescent reporter strains in pairs of sister cells in the following cell generation (AB64 stage--when there are 32 ABa descendants). scRNAseq data are consistent with mRNA being expressed in the parent cell, with reporter detection presumably delayed by the time required for the fluorophores of mCherry or GFP to mature (Figure S5). This raises the question of whether *ref-2* expression requires the ABa-specific transcription factors *tbx-37* and *tbx-38*. These genes encode redundant paralogous T-box family transcription factors which are expressed throughout the ABa lineage and are required for multiple cell fate decisions within ABa [46,50,55]. Confirming previous reports [46,50], a TBX-38::GFP knock-in reporter made by CRISPR [50] is expressed throughout the ABa lineage and is detectable at least from the AB16 to AB128 stages (Figure S10) (*tbx-37* and *tbx-38* RNA are detectable by the AB4 stage [56]). To test whether *tbx-37* and *tbx-38* are necessary for *ref-2* expression, we measured expression of the *ref-2* promoter reporter in embryos carrying homozygous deletions of both *tbx-37* and *tbx-38*. We found that in the absence of *tbx-37* and *tbx-38,* nearly all *ref-2* promoter expression in the ABa lineage is lost (Figure 5H; Figure S11), while expression in the MS lineage was maintained. Thus, *tbx-37* and *tbx-38* are necessary for the anterior-biased expression of *ref-2* in ABa.

### The −3.9 kb enhancer contains three non-overlapping sub-enhancers that independently drive anterior expression in the ABa lineage

Since the *ref-2* −3.9 kb enhancer drives expression in a similar pattern to the early expression of the full promoter and protein reporters, and its deletion from the *ref-2* promoter results in a reduction of anterior-biased expression in portions of the ABa lineage, it is a primary enhancer responsible for driving the early anterior-biased expression of *ref-2*. To map features necessary or sufficient for anterior expression, we tested the activity of eight truncated versions of the *ref-2* - 3.9 kb enhancer to identify minimal regions of the enhancer sufficient to drive anterior expression in the ABa lineage (Figure 6, Figure S12). Instead of a single minimal region, we found three non-overlapping regions within the −3.9 kb enhancer that each are sufficient to drive anterior-biased enhancer activity in the ABa lineage: a promoter-distal region (bases 1 to 200), a medial region (bases 227 to 304) and a promoter-proximal region (bases 320 to 449) (Figure 6B, C). Of these regions, the proximal region drives the most consistent expression and this expression is most similar to the full length −3.9 kb enhancer construct (Figure 6D), whereas the medial and distal fragments drive more variable expression and in a subset of these lineages. Thus, although integration of information between these sub-enhancers is likely necessary for robustness of expression pattern, the 130 bp proximal fragment provides a model system for further dissection to identify anterior expression regulators.

**Figure 6:**
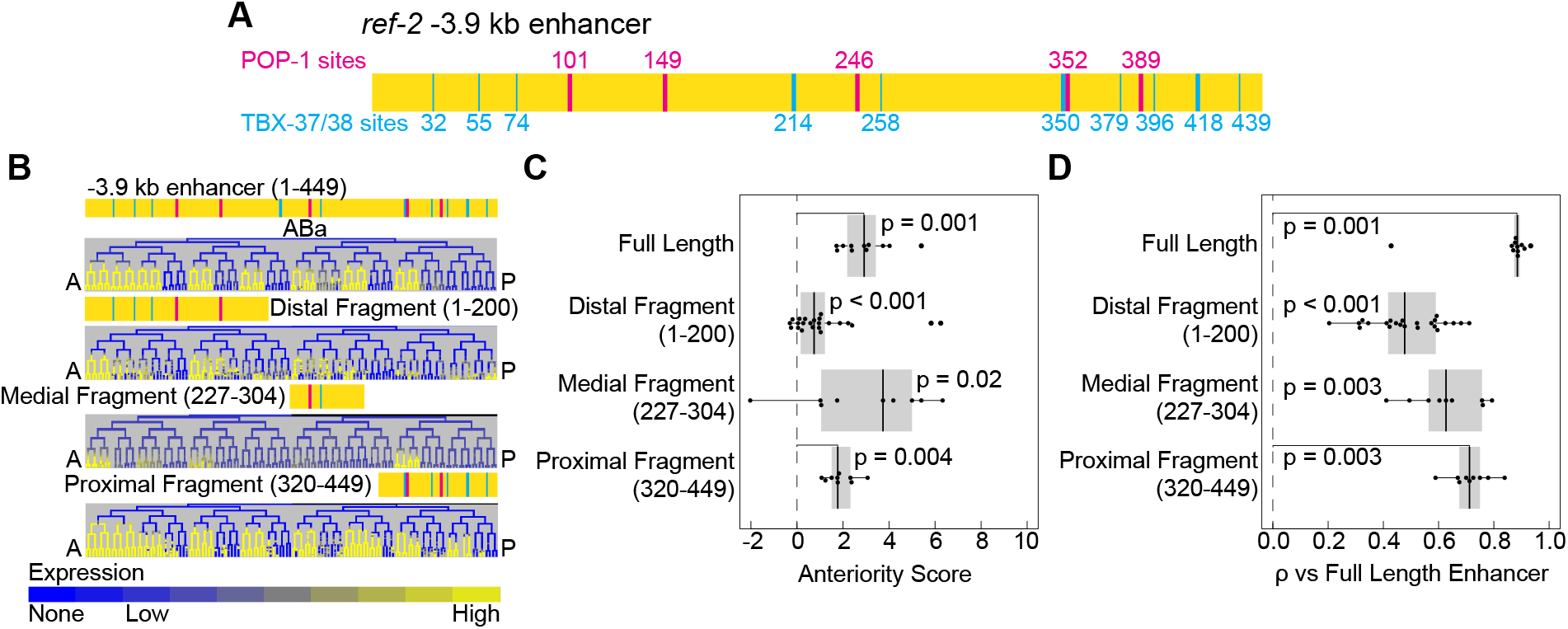
Three non-overlapping fragments of the −3.9 kb enhancer are each sufficient to drive anterior-biased expression in the ABa lineage of the early embryo. A) Model of the *ref-2* −3.9 kb enhancer with predicted TBX-37/38 sites (cyan, predicted high affinity sites indicated by thick line and predicted low-affinity by a thin line), and predicted high affinity POP-1 sites (magenta). B) Expression patterns driven by full-length *ref-2* −3.9 kb enhancer and by minimal fragments. C-D) Box plots displaying the anteriority scores (C) and Spearman’s ρ (D) for the full-length *ref-2* −3.9 kb enhancer and minimal fragments. Lineages used to determine anteriority scores are the same as in Figure 5J. Spearman’s ρ analysis uses the full ABa lineage and was calculated relative to the mean expression of the full-length −3.9 kb enhancer. A complete set of enhancer truncations tested is displayed in Figure S12.

### TBX-37/38 binding sites are required for anterior expression of the proximal region

The proximal fragment exhibits broad sequence conservation with related nematodes, suggesting it contains multiple important TF binding sites (Figure 5A). To identify sequences necessary for anterior expression, we scrambled each 15 base-pair region along the length of the enhancer fragment and tested the resulting sequences for enhancer activity (Figure S13B, C, D). Scrambling a single region comprising base pairs 410-424 (coordinates relative to the full-length enhancer) resulted in a complete loss of the anterior-biased expression in the ABa lineage, indicating that this region is necessary for this expression (Figure 7B; Figure S13B). Scrambling the most proximal regions 425-439 and 435-449 resulted in a loss of expression in subsets of the ABa lineage (in ABar and ABalp) but maintained anterior expression in the ABala lineage (Figure S13B). Expression was largely maintained when other regions were mutated, suggesting sequences in these regions are not individually necessary for anterior-specific expression despite their conservation.

**Figure 7:**
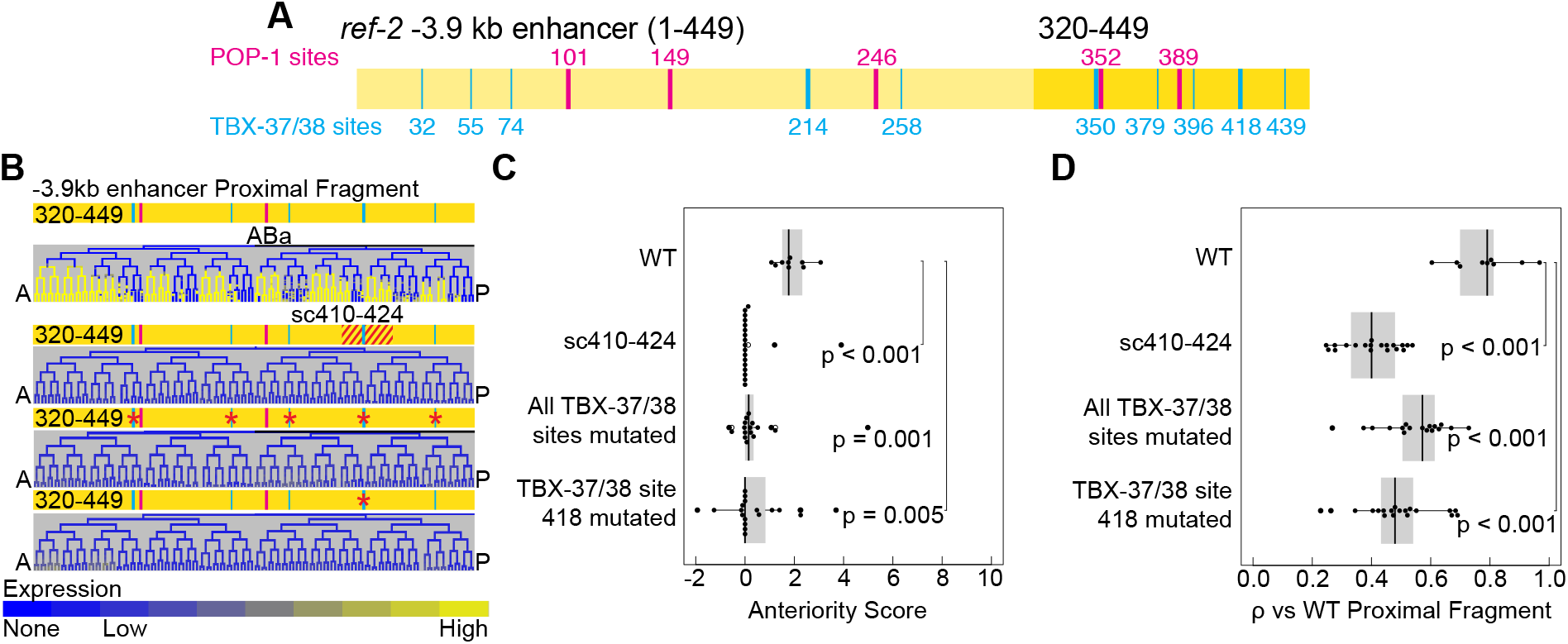
The ABa-expressed transcription factors *tbx-37* and *tbx-38* are required for the anterior-biased expression of *ref-2* in the ABa lineage. A) Model of the *ref-2* −3.9 kb enhancer with proximal fragment (base pairs 320-449) highlighted. Predicted TBX-37/38 and POP-1 sites are indicated as in Figure 6. B) Expression patterns driven by the proximal fragment of the *ref-2* - 3.9 kb enhancer with its WT sequence, with base pairs 410-424 scrambled, with all predicted TBX-37/38 sites mutated, and with *TBX_418_* mutated. Expression driven by the proximal fragment with additional regions scrambled or with two predicted POP-1 sites mutated are displayed in Figure S13. C-D) Anteriority scores (C) and Spearman’s ρ (D) for the proximal fragment with its WT sequence, with base pairs 410-424 scrambled, with all predicted TBX-37/38 sites mutated, and with *TBX_418_* mutated. Lineages used to determine anteriority scores are the same as in Figure 5J. Spearman’s ρ analysis uses the full ABa lineage and was calculated relative to the mean expression of the WT −3.9 kb enhancer proximal fragment.

The proximal region includes five predicted TBX-37/38 sites, two of which are predicted to have high affinity for TBX-37 and TBX-38. Two of the sites overlap with either the 410-424 region that is essential for ABa-specific expression or the 425-449 region that is required for a subset of ABa-specific expression (Figure 7A, B; Figure S13A, B). To determine whether this region is bound by TBX-37 and TBX-38 *in vivo,* we mined a recently published dataset that measured genome-wide TBX-37 and TBX-38 binding by ChIP-seq. Both factors show binding in the proximal region of the −3.9 kb enhancer (Figure 5A) [50].

To determine whether the TBX-37/38 sites are required for anterior expression, we measured the activity of the proximal fragment after mutating the two central nucleotides of all TBX-37/38 sites. The mutated enhancer fragment did not drive anterior-specific expression in the ABa lineage (Figure 7B, C, D; Figure S13B, C, D; Figure S14), confirming the importance of these sites. Additionally, the loss of anterior-biased expression following the mutation of these sites indicates that the two predicted high affinity POP-1 sites in this fragment are not sufficient to drive anterior-biased expression. Also, these sites are not necessary for the anterior-biased expression driven by this enhancer fragment, as the loss of these sites does not result in a loss of the anterior-biased expression (Figure S13B, C, D). In fact, concatemers of POP-1 sites are sufficient to drive posterior-biased expression (Figure S13F) [13,53].

Since the high affinity site beginning at position 418 (*TBX_418_*) overlaps the region (410-424) required for expression, we hypothesized that this specific TBX-37/38 site may be required for anterior expression in the ABa lineage. To test this, we tested the activity of the proximal fragment with only the central two nucleotides of *TBX_418_* mutated (Figure 7B, Figure S13B, E). This mutation resulted in a loss of robust anterior-biased expression driven by the proximal fragment. Therefore, TBX-37 and TBX-38 bind to the *ref-2* −3.9 kb enhancer proximal fragment and *TBX_418_* is necessary for it to drive robust anterior-biased expression (Figure 5A; Figure 7B, C, D; Figure S13B, C, D; Figure S14), suggesting that TBX-37 and TBX-38 directly regulate *ref-2* expression.

### Concatemerized TBX sites are sufficient to drive anterior-specific expression in the ABa lineage

Given that *tbx-37* and *tbx-38* are expressed in both anterior and posterior cells within the ABa lineage, we hypothesized that site *TBX_418_* would be sufficient to drive broad expression in ABa, with other sequences in the enhancer required for anterior specificity. To test this, we tested the activity of synthetic enhancers comprising concatemerized copies of *TBX_418_* separated by either short (6-7 bp) or long (11-15 bp) flanking sequences from its endogenous context (Figure 8A). Surprisingly, both of these reporters drove anterior-specific expression in the ABa lineage similar to that driven by the full proximal fragment (Figure 8B, C, D). This indicates that either *TBX_418_* or its flanking sequences are sufficient both for ABa lineage expression, and for anterior-specific expression within this lineage. To test the possibility that the flanking sequences are responsible for anterior-specificity, we scrambled those sequences while leaving the central *TBX_418_* site intact in the context of the long-flanking-sequence construct. The resulting reporter was again expressed only in anterior sublineages similar to the full proximal fragment (Figure 8A, B, C, D). Thus, when multimerized, *TBX_418_* is sufficient both for ABa expression and for anterior specificity without additional non-overlapping sequences.

**Figure 8:**
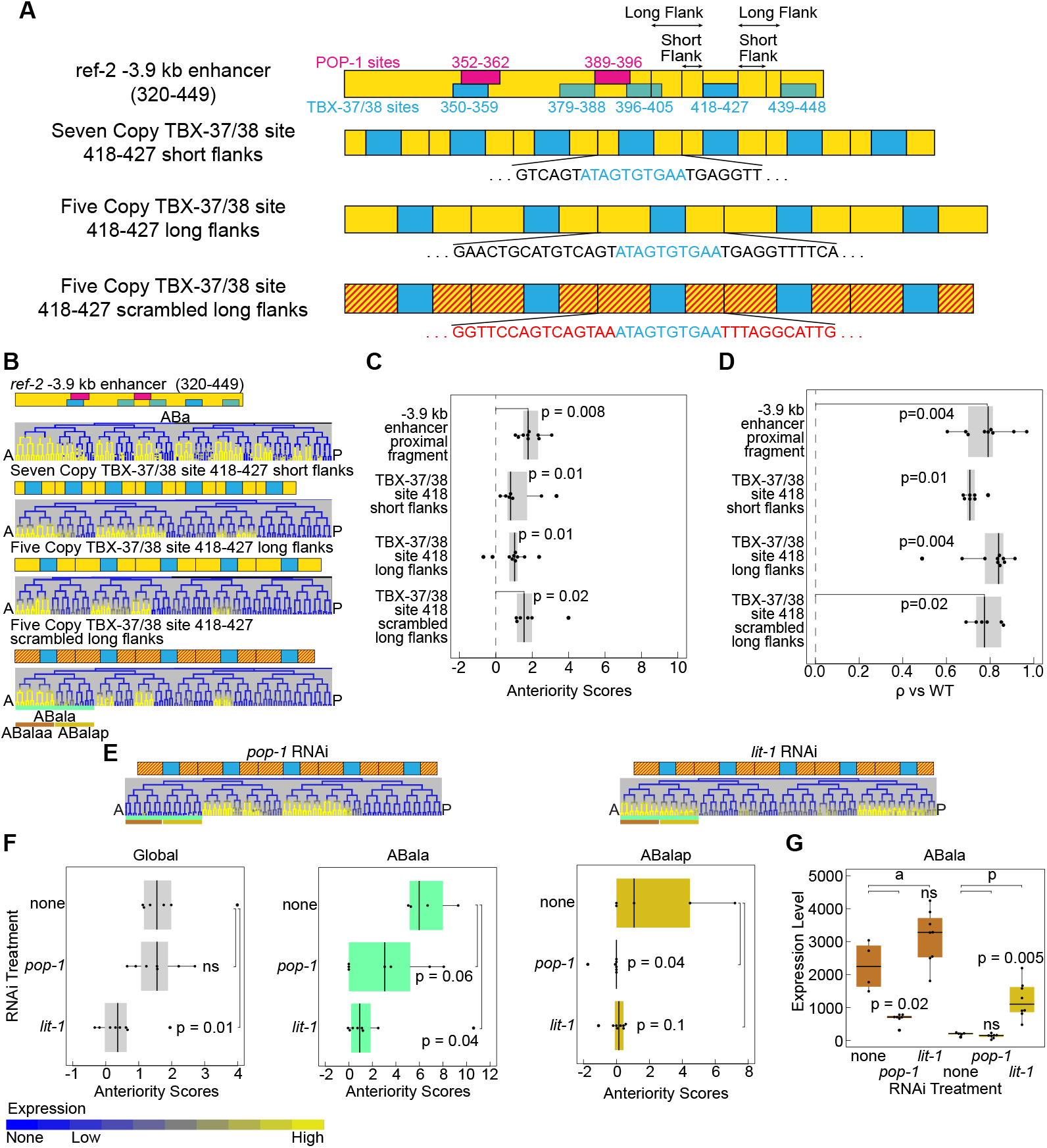
*TBX_418_* in the *ref-2* −3.9 kb enhancer is sufficient when concatemerized to drive anterior-biased expression in the ABa lineage. A) Models of the proximal fragment of the *ref-2* −3.9 kb enhancer, the *TBX_418_* seven-site concatemer with 6-7 base pair flanking regions, the *TBX_418_* five-site concatemer with 11-15 base pair flanking regions, and the *TBX_418_* five-site concatemer with 11-15 base pair scrambled flanking regions. B) Expression patterns driven by the proximal fragment of the *ref-2* −3.9 kb enhancer and each TBX-37/38 site concatemer. C-D) Anteriority scores (C) and Spearman’s ρ (D) for the proximal fragment of the *ref-2* −3.9 kb enhancer and each TBX-37/38 binding site concatemer. E) Expression patterns driven by the *TBX_418_* five-site concatemer with 11-15 base pair scrambled flanking regions after *pop-1* RNAi or *lit-1* RNAi. F) Anteriority scores for the *TBX_418_* five-site concatemer with 11-15 base pair scrambled flanking regions in indicated lineages following indicated RNAi treatments. G) Quantification of reporter expression for the *TBX_418_* five-site concatemer with 11-15 base pair scrambled flanking regions in the ABala lineage. Expression is shown under WT conditions and following *pop-1* or *lit-1* RNAi. Lineages used to determine global anteriority scores are the same as in Figure 5J. Spearman’s ρ analysis uses the full ABa lineage expression pattern and was calculated relative to the WT proximal fragment.

### Anteriorly-biased expression of concatemerized TBX sites is regulated by *pop-1*

*ref-2* anterior-specific expression is dependent on *pop-1* and *sys-1*, and a concatemerized TBX-37/38 binding site recapitulates part of this expression. This raises the question of whether the TBX-37/38 binding site expression is *pop-1-*dependent. To test this, we examined the expression of the *TBX_418_* concatemer with scrambled flanking regions after *pop-1* RNAi. Knockdown of *pop-1* results in changes in the expression pattern, including a significant decrease in anterior expression bias in the ABalap lineage and a significant decrease in expression level in the ABalaa lineage (Figure 8E, F, G). We also tested the expression effects of *lit-1* RNAi, which causes an increase in nuclear POP-1 levels in posterior cells [16,57,58]. Knockdown of *lit-1* results in a significant decrease in the anterior bias of the *TBX_418_* concatemer expression across several sublineages of ABa, including a significant increase in expression in the ABalap lineage (Figure 8E, F, G). Some of these changes mirror the fate transformations known to occur after *pop-1* or *lit-1* RNAi, but others, including the increase in ABalap expression after *lit-1* RNAi, do not [14,59]. Thus, our results indicate that either POP-1 or its targets influence both the lineage specificity and anterior specificity of expression driven by *TBX_418_*.

## DISCUSSION

Previous work has definitively shown that the Wnt pathway components POP-1 and SYS-1 regulate posteriorly expressed genes in the *C. elegans* embryo via the canonical Wnt asymmetry pathway [12,13,36]. What has been less clear is how the similar number of anteriorly expressed genes are regulated. We have demonstrated that a set of anteriorly expressed genes, *tbx-35*, *ceh-51*, *elt-1*, *irx-1*, and *ref-2*, require POP-1 and SYS-1 for their anterior-biased expression. Therefore, POP-1 and SYS-1 regulate the expression of anterior-biased genes. The mechanism by which POP-1 and SYS-1 mediate this regulation remains to be determined. Previous work demonstrated that posterior genes can be regulated by POP-1 binding directly to POP-1 binding sites in their upstream regulatory regions [53]. Thus, a simple model of POP-1 regulation of anterior genes would be indirect, with POP-1 activating the expression of repressors in posterior cells. For example, *pop-1* RNAi is known to cause the anterior MS cell to adopt the fate of its posterior sister E due in part to derepression in MS of the E-specific classic Wnt target *end-1*. *end-1* (or its targets) could then act to repress expression of MS-specific genes. This mechanism would predict that anterior genes would be expressed later than posterior genes, or that anterior genes would be expressed early in both anterior and posterior cells, but maintained only in anterior cells. This can happen. For example, in the C lineage *elt-1* is expressed in both anterior and posterior sister cells at the 4C stage, but is only maintained in the anterior granddaughters of C (Caa and Cpa), perhaps due to its repression by the Wnt target *hlh-1* in the descendants of the posterior granddaughters of C (Cap and Cpp) [33]. What we observe in most cases, however, is that anterior gene expression is activated at about the same time as posterior gene expression in sister lineages (Figures S1, S2, S3, S4, S5, and S6, Zacharias et al. 2015 [13]). This observation suggests that POP-1 could regulate some anterior genes directly. Some *pop-1* dependence of anterior genes certainly could be explained by upstream anterior-to-posterior cell fate conversions following *pop-1* RNAi; however, our results as a whole cannot be explained by the known pattern of fate transformations [46].

Indirect *pop-1*-dependent regulation of anterior genes does not explain all of its role in anterior gene expression regulation. For instance, previous work showed that loss of muscle-specifying TFs (MRFs) expressed in Cap and Cpp (*hlh-1*, *unc-120*, and *hnd-1*) is not sufficient to permit ectopic expression of *elt-1* in these cells or to convert them to an epidermal fate [33,60]. Therefore, another factor, perhaps *pop-1* acting directly, must be responsible for restricting *elt-1* to the anterior C granddaughters. Our results identified a requirement for *pop-1* in anterior expression of *elt-1* but not posterior repression in the C lineage. The proposed role of the MRFs in repressing *elt-1* expression [33,60,61] suggests that this effect could partly be due to misexpression of MRFs in the anterior C granddaughters, which could be tested in the future.

Direct activation of genes by POP-1/TCF when Wnt is absent occurs in other systems. For example, in *Drosophila* TCF/Pangolin can act in a reverse transcriptional switch such that when it is bound directly to DNA at non-optimal sites it can activate transcription when unbound by β-catenin, and repress transcription when bound to β-catenin. This difference in the regulatory activity of TCF is mediated by an allosteric conformational change in the structure of TCF [22]. In *C. elegans*, POP-1 bound together with REF-2 at the *ttx-3 cis-*regulatory region, may similarly undergo allosteric changes, leading to the opposite regulation of target genes, such that POP-1 activates the expression of *ttx-3* when bound with REF-2 in the absence of SYS-1 and represses the expression of *ttx-3* when bound with REF-2 and SYS-1 [20,21]. Our findings are consistent with a similar mechanism regulating *ref-2* itself.

Our observation that five copies of a single 10 base-pair TBX-37/38 binding site are sufficient to drive anterior-biased expression in the ABa lineage suggests that a POP-1 site is likely unnecessary for anterior expression. However, *tbx-37* and *tbx-38* cannot explain this anterior bias alone, as these factors are expressed throughout the ABa lineage. We showed that *pop-1* is required for both anterior expression and posterior repression of the *ref-2* promoter and for anterior expression of a TBX-37/38 binding site concatemer. If this role of POP-1 is direct, it likely acts by binding to other factors, possibly TBX-37 and TBX-38, rather than to DNA directly since the concatemer contains no sequences with high affinity for POP-1. Consistent with this, TBX-38 and POP-1 were previously found to physically interact with each other in a yeast 2-hybrid assay [62]. Intriguingly, several other T-box factors are expressed early in the AB lineage [50,56]; future work should determine if these play a role in restricting the activity of these sites to anterior lineages. For instance, *TBX_418_* is predicted to be bound by several other *C. elegans* T-box factors. Other T-box factors expressed in the early ABa lineage could compete for binding at this site, with POP-1 altering the relative activity of these factors between anterior and posterior cells. Additionally, it is unclear whether *tbx-37* and *tbx-38* also regulate posterior gene expression in the ABa lineage; intriguingly, the expression of *pha-4*, another anterior gene, in the ABa lineage is also dependent on *tbx-37* and *tbx-38* [46].

In contrast with our observation that *TBX_418_* concatemers are sufficient to drive anterior-biased expression, Murgan et al. 2015 found that a REF-2 binding site concatemer is not sufficient to drive expression in the anterior AIY mother cell. Instead, Murgan et al. identified binding sites for helix-loop-helix family transcription factors that are sufficient to drive expression in the AIY mother and its posterior sister cell, and that combining these with REF-2 binding sites restricts expression to the AIY mother. Thus, REF-2 acting through ZIC binding sites is not sufficient to activate expression without cooperating factors binding other sites whereas TBX-37 and TBX-38 may be sufficient to do so. Other factors could also play a role, for example we used a *pes-10* minimal promoter, whereas Murgan et al. did not include a minimal promoter in their constructs, and there could be differences in sensitivity between the microscopy techniques [20].

Classic work in many species, including flies and vertebrates, has found that developmental genes are often regulated both by partially redundant enhancers and modular enhancers that are responsible for regulating distinct portions of the genes’ expression patterns [63,64]. The extent to which *C. elegans* genes are regulated by such distal enhancers vs promoter proximal elements has been unclear. We found that the predicted enhancers in the promoter region of *ref-2* function modularly, such that each enhancer drives different expression patterns, presumably providing multiple inputs to fine-tune the expression pattern of *ref-2*. Combined with other recent studies this adds to evidence that enhancer-mediated regulation is widespread during *C. elegans* embryonic development [54,65–69]. Additionally, we observe modularity within the −3.9 kb enhancer, with three non-overlapping regions of the enhancer each sufficient to drive anterior-biased expression. Since the protein reporter is expressed in more lineages than the promoter reporter, there are likely even more enhancers outside the promoter region that regulate expression of *ref-2*.

Several studies have identified evidence for multiple functions of *ref-2*. *ref-2* was first identified as being required for the production of the Pn.p ventral epidermal cells and for the inhibition of their fusion to the epidermal syncytium hyp7 [39]. *ref-2* is also required for the initiation of the differentiation of the cholinergic neuron AIY [40]. Also, *ref-2* has been found to be required for female fate during sexual development [70]. We have demonstrated a role for *ref-2* in embryonic development. *ref-2* is required for robust WT cell positioning and cell division timing in anterior lineages. Intriguingly, an insect ortholog of *ref-2*, *odd-paired*, acts as a regulator of anterior-posterior expression and is an anterior determinant in several species [71,72], suggesting the function of *ref-2* in anterior fate regulation may be ancestral. Because the defects in cell position and division timing are only partially penetrant, there are likely other, partially redundant regulators of cell position and cell division timing that act with the sole *C. elegans* ZIC homolog *ref-2* during embryonic development. Further experiments, including suppressor and enhancer screens and co-mutation of other transcription factors will need to be performed to identify other factors that regulate cell position and division timing in conjunction with *ref-2*.

## METHODS

### *C. elegans* culture and strain generation

*C. elegans* strains (Table S1) were maintained at standard growth temperatures on OP50 *E. coli* on NGM plates (Table S2). RNAi knockdown was performed by feeding, as previously described [73,74]. We validated the efficacy of *pop-1* RNAi by measuring the cell cycle delay resulting from the transformation of the anterior MS lineage into an E-like lineage, the efficacy of *sys-1* RNAi by failure of morphogenesis and embryonic death, and the efficacy of *lit-1* RNAi by measuring the cell cycle shortening resulting from the transformation of the posterior E lineage into an MS-like lineage (Table S2) [13,34,59]. Enhancer reporter strains were generated by microinjection into RW10029, the GFP histone strain used for lineage tracing. Injection cocktails used to create extrachromosomal array transgenes consisted of a linear reporter DNA construct at 10 ng/µL, with 5 ng/µL *myo-2*p::GFP, and 135 ng/µL pBluescript vector (or highest concentration possible whenever 135 ng/µL was impossible to make) and were injected using a Narishige MN-151 micromanipulator with Tritech microinjector system. Enhancer reporters were maintained in extrachromosomal arrays by screening for worms expressing the *myo-2*p::GFP reporter, which have a green fluorescent pharynx. Most enhancer reporters, including all for which loss of expression was observed, were analyzed in at least two strains to control for differences in copy number and extrachromosomal array artifacts. Other strains were created through crosses using standard approaches (Table S1).

### Generation of transgenes

Candidate enhancers were amplified by PCR from *C. elegans* N2 strain genomic DNA with Phusion HF polymerase (New England Biosciences) and gel purified (Qiagen). Enhancer fragments, mutated enhancers, and binding site concatemers were ordered as either gBlocks or Ultramers from Integrated DNA Technologies (Coralville, Iowa) and were amplified with Phusion HF polymerase (New England Biosciences) with overhangs for stitching and either gel or PCR purified (Qiagen). Enhancer reporters were produced by fusing via PCR stitching these constructs to a *pes-10* minimal promoter::HIS-24::mCherry::let-858 3’UTR fragment amplified from the POPTOP plasmid [53] (Addgene #34848). The enhancer reporters were purified with a PureLink PCR purification kit (ThermoFisher) and/or gel purified. The desired product was determined by its size based on gel electrophoresis. We identified putative transcription factor binding sites using CIS-BP (http://cisbp.ccbr.utoronto.ca) [75,76].To mutate the TBX-37/38 sites, the central two nucleotides were altered (e.g. ATAG**<u>TG</u>**TGAA changed to ATAG**<u>GT</u>**TGAA for *TBX_418_*). For the construct with all TBX-37/38 sites mutated, after mutating the five primary sites, subsequent analysis revealed two weaker sites. All of these predicted TBX-37/38 sites were mutated. The reporter construct for the *ref-2* promoter lacking the −3.9 kb enhancer was amplified from N2 genomic DNA by PCR and PCR stitched to the *pes-10* minimal promoter::HIS-24::mCherry reporter, and, thus, drives expression through the *pes-10* minimal promoter. The *pes-10* promoter drives no consistent embryonic expression on its own prior to elongation [13,53,54] and is compatible with a wide variety of enhancers [13,53] (Table S5).

### Quantitative comparisons

All quantitative comparisons were performed using R version 4.0.3 (The R Foundation for Statistical Computing), Microsoft Excel version 16.16.27, or Python version 2.7.16. For the main figures, a minimum N of 6 was used for all experiments, except for ELT-1::GFP expression after *sys-1* RNAi, for which N = 3. For the supplemental figures, a minimum N of 2 was used (largely for perturbations which appeared similar to wild-type at this level). Code and raw data for these analyses are available at https://github.com/jisaacmurray/ref-2_paper.

### Expression and phenotypic analysis by 4D imaging

We obtained confocal micrographs using a Leica TCS SP5 or Stellaris scanning confocal microscope (67 z planes at 0.5 µm spacing and 1.5 minute time spacing, with laser power increasing by 4-fold through the embryo depth to account for attenuation of signal with depth). Embryos obtained from self-fertilized hermaphrodites were mounted in egg buffer/methyl cellulose with 20µm beads used as spacers and imaged at 22°C using a stage temperature controller (Brook Industries, Lake Villa, IL) [77]. We used StarryNite software to automatically annotate nuclei and trace lineages [29]. We corrected errors from the automated analysis and quantified reporter expression in each nucleus relative to the local background (using the “blot” background correction technique) with AceTree software as previously described (Table S3) [23,25].

### Anterior gene WT to RNAi within lineage comparisons

To compare the expression levels of the anterior genes *tbx-35*, *ceh-51*, ELT-1, *irx-1*, and *ref-2* and the *TBX_418_* concatemer with 11-15 base pair scrambled flanking regions between WT and RNAi embryos within the expressing lineages and their posterior sisters, we calculated for each embryo the mean reporter expression across all measurements within each lineage (all descendant cells and time points) starting with the cell stage that expresses the reporter at the time cells are born or shortly afterwards to the last time point in the indicated lineage. For expression level comparisons of extrachromosomal array transgenes, we normalized expression values to the 75th percentile expression value of the strain with the higher 75th percentile expression value across all measured cells in the indicated lineage within each RNAi treatment. The expression analyzed for the MS lineage starts at an earlier stage than that for other lineages, since following *pop-1* RNAi the cell divisions are so dramatically delayed that reporter expression comes on in earlier cell generations in these embryos. The cell generations used in each anterior gene comparison are noted in Table S4.

### Anteriority Scores

To measure the magnitude of anterior expression bias, we developed a measure that we refer to as the anteriority score. To determine the anteriority scores, first we chose a minimum expression cutoff and set all values below this cutoff to 0 and subtracted the minimum cutoff from all expression values greater than or equal to the cutoff (an expression value of 200 was used as the cutoff for all anteriority score analyses, except a value of zero was used for the TBX-38::GFP reporter, which has very low fluorescence intensity compared to the other reporters analyzed). Next, we averaged the expression levels of the reporter of interest in the cells of each anterior and each posterior sister lineage of interest. For the *ref-2* promoter and enhancer constructs, as well as the TBX-37/38 site concatemers, we analyzed anterior and posterior lineages descended from the cells ABala, ABalaa, ABalap, ABalpa ABalpp ABaraa, ABarap, ABarpa, and ABarpp. We used expression values starting at the generation with ABaxxxxxx cells (AB128 stage) until the time point of the last division of AB128 stage cells in the ABa lineage. Next, we calculated the means of the average expression levels of all of the anterior lineages and all of the posterior lineages analyzed for each embryo. We then added a pseudocount of 1 to each of the anterior and posterior embryo means. For each experimental group, the adjusted average anterior expression is divided by the adjusted average posterior expression, and the base 2 logarithm of the result is reported--Anteriority Score = log_2_((anterior mean + 1)/(posterior mean +1)).

### Correlation analysis

Correlation analysis was done by Spearman’s ρ, comparing each embryo’s expression to the average expression of the control group. Spearman’s ρ was calculated in R using the “cor” function. For Spearman’s ρ we used all the data from the lineages of interest without excluding any cell generations or having a minimum value cutoff. We included cells born before the onset of expression here to include information about the cell generation of expression onset in the analysis. For the analysis comparing the *ref-2* promoter with its enhancers we used the full somatic lineage starting with the blastomeres AB, EMS, C, and D. For all other analyses we used only the ABa lineage.

### Mutant cell position and cell division analysis

We identified cell position and cell cycle defects as previously described [41]. We corrected for differences in global division rates, and considered divisions as defective if they deviated from the wild-type cell cycle length by at least five minutes and had a z-score greater than three. Cell positions were corrected for differences in embryo size and rotation, and considered defective if they deviated from the expected wild type position by at least five microns, had a z-score greater than five, and a nearest neighbor score greater than 0.8.

### Statistical analyses

Statistical significance of differences in anterior or posterior expression between untreated and RNAi-treated embryos was tested using the two-tailed Wilcoxon Ranked Sum test. Statistical significance of differences in Spearman’s ρ or Anteriority Score were tested using the one-tailed Wilcoxon Ranked Sum test to test for differences from either the control group or from 0, as indicated in each figure. The p-values calculated by the Wilcoxon Ranked Sum test were corrected for multiple comparisons using the false discovery rate method in R. Significance of overlap between positional or cell division defects and expression status were assessed using a Chi-squared test.

---

**Figure S1:**
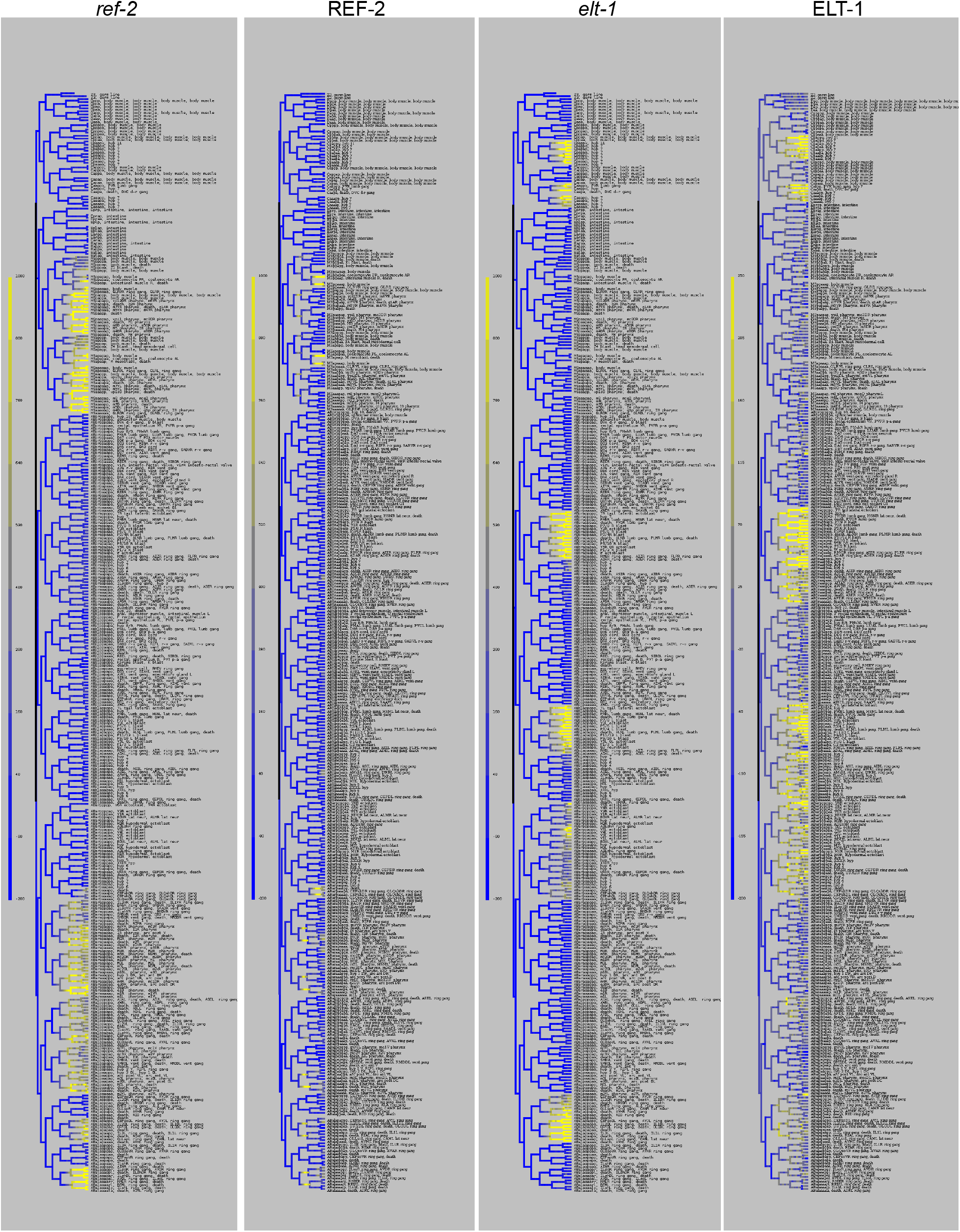
Several early embryonic genes are expressed with anterior bias. Full lineage expression patterns are shown for integrated transgenic promoter reporters of *ref-2* and *elt-1*; and for integrated transgenic protein reporters of REF-2 and ELT-1.

**Figure S2:**
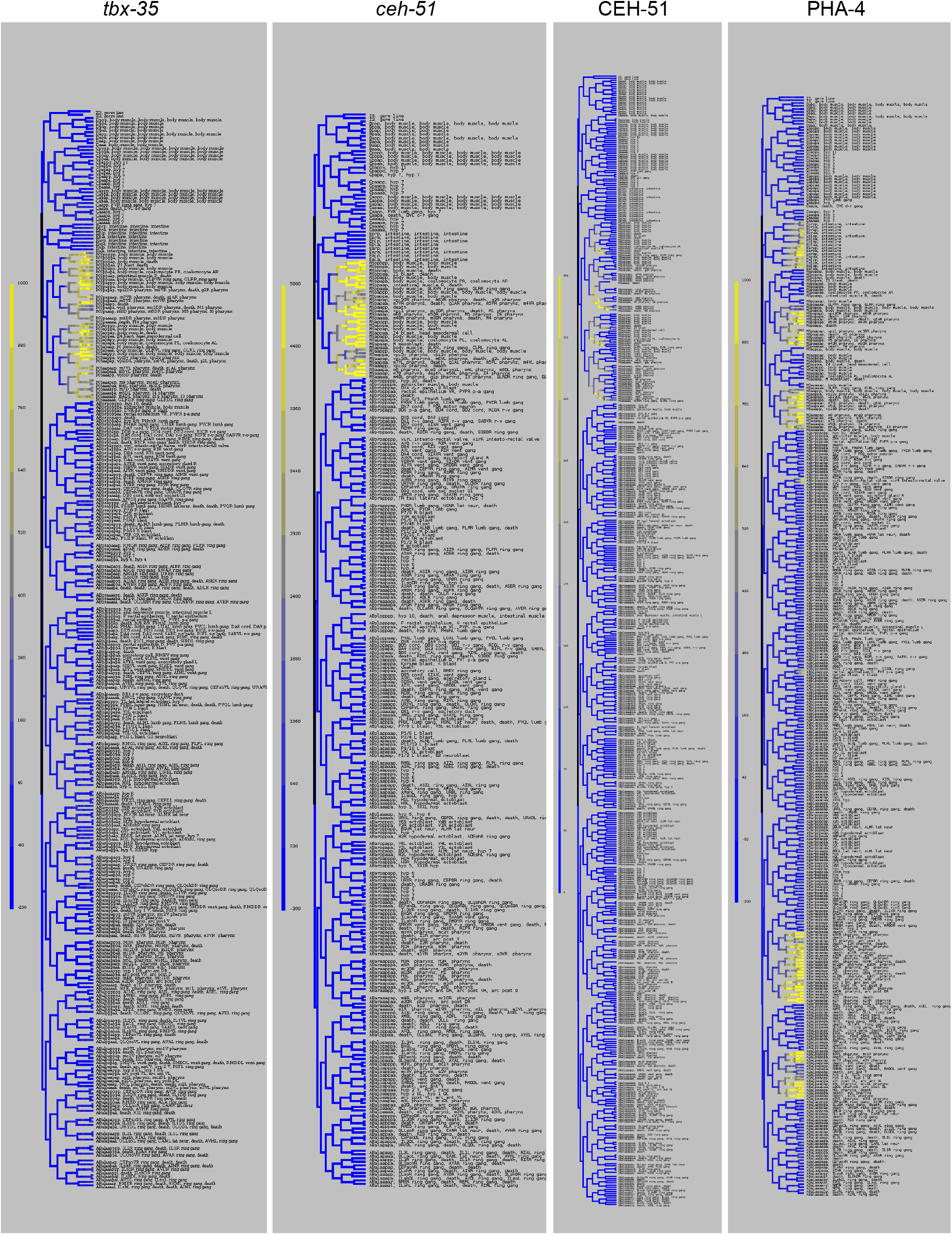
Several early embryonic genes are expressed with anterior bias. Full lineage expression patterns are shown for integrated transgenic promoter reporters of *tbx-35* and *ceh-51*; for an integrated transgenic protein reporter of PHA-4; and for a CRISPR knock-in protein reporter of CEH-51.

**Figure S3:**
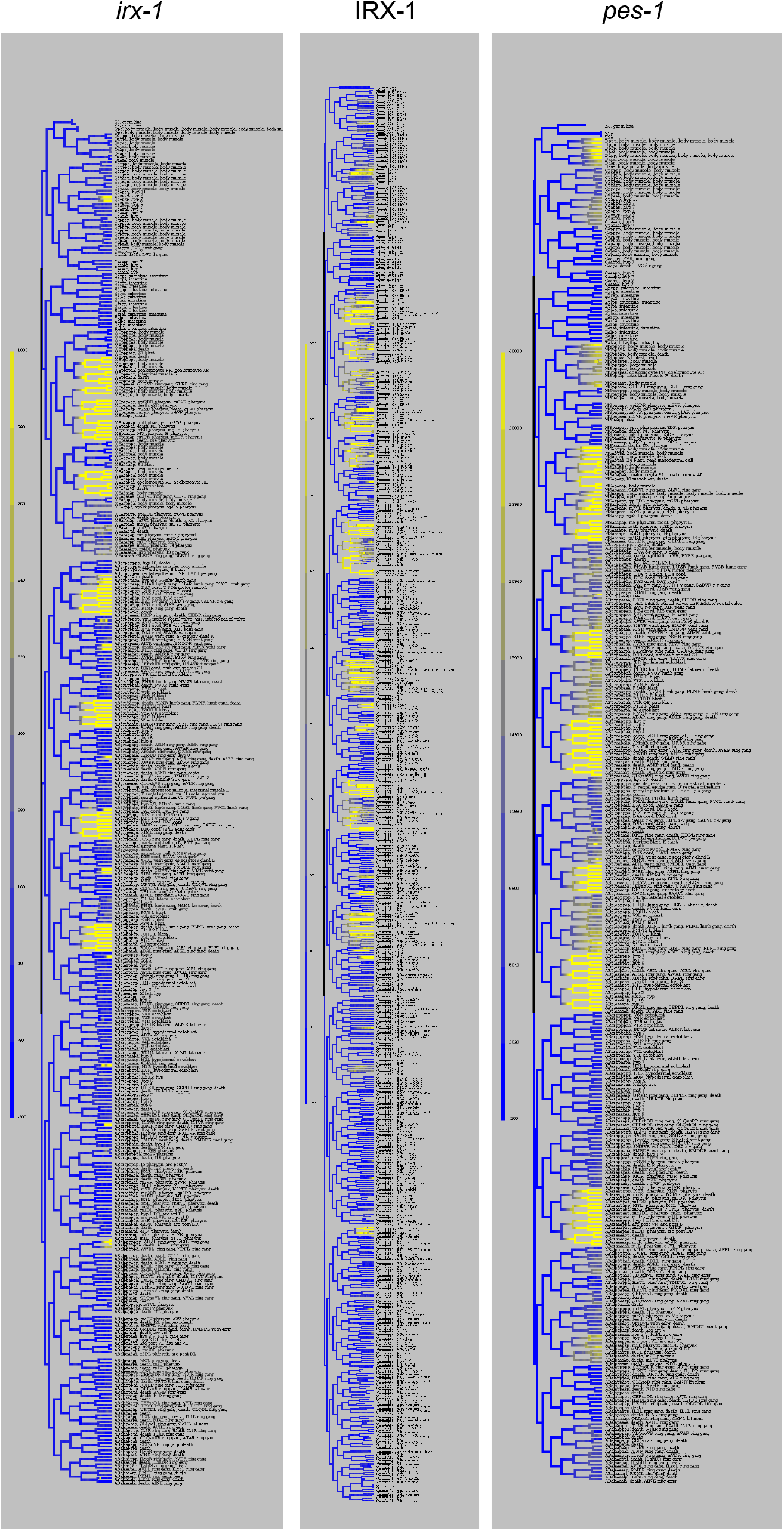
Several early embryonic genes are expressed with anterior bias. Full lineage expression patterns are shown for integrated transgenic promoter reporters of *irx-1* and *pes-1*; and for an integrated transgenic protein reporter of IRX-1.

**Figure S4:**
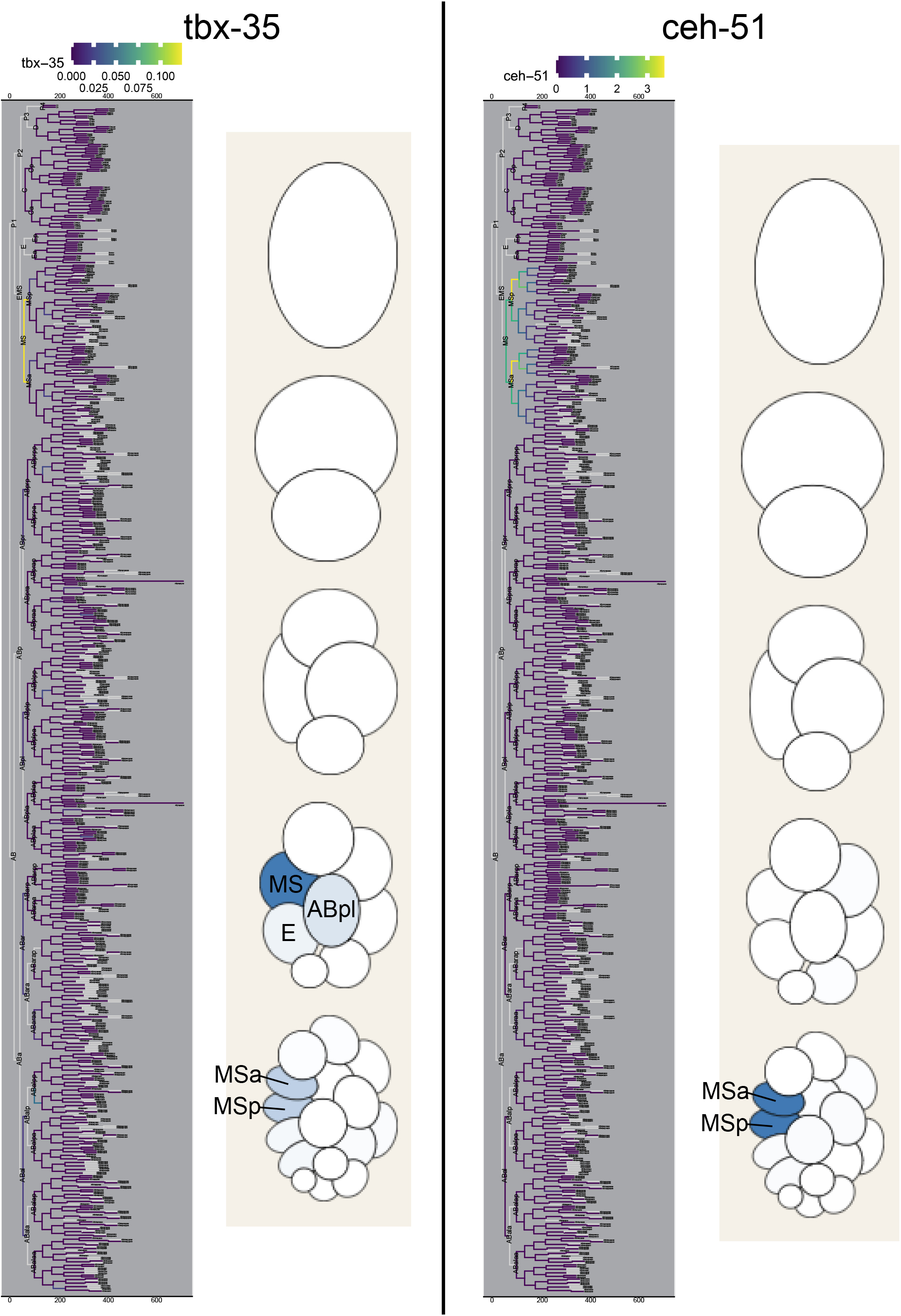
scRNA-seq data confirm imaging-based expression data. scRNA-seq expression data for *tbx-35* and *ceh-51*. *tbx-35* expression lineage derived from Packer et al. 2019 indicates expression in the daughters of MS (MSa and MSp) [32]. Early *C. elegans* embryo diagram derived from Tintori et al. 2016 indicates *tbx-35* expression in MS and its daughters, as well as low-level expression in E and ABpl [56]. *ceh-51* expression lineage derived from Packer et al. 2019 indicates expression in the descendants of MS from the MS2 through the MS16 generation (MS daughters through MS great-great granddaughters) [32]. Early *C. elegans* embryo diagram derived from Tintori et al. 2016 indicates *ceh-51* expression in the daughters of MS [56]. In both cases scRNA-seq expression data agree with imaging-based expression data, albeit with expression being detected earlier and persisting for fewer cell generations than for the fluorescent reporters (Figure 3 and Figure S6). Expressing cells in early embryo diagrams are labeled with their names. Gray branches on lineage trees indicate cells for which distinct transcriptomes could not be determined.

**Figure S5:**
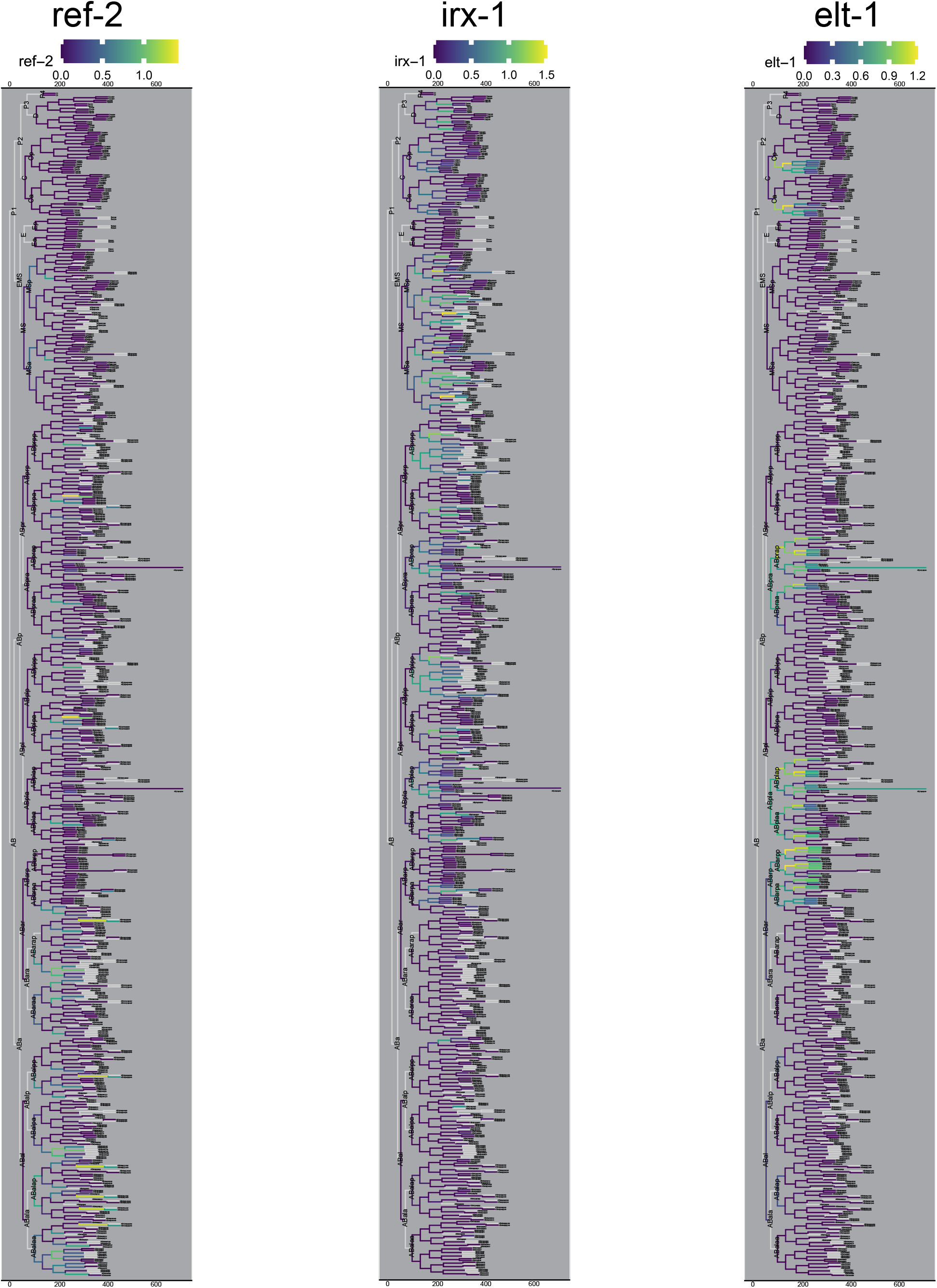
scRNA-seq data confirm imaging-based expression data. scRNA-seq expression data for *ref-2*, *irx-1*, and *elt-1* derived from Packer et al. 2019 [32]. scRNA-seq expression data largely agree with imaging-based expression data, albeit with expression being detected earlier and persisting for fewer cell generations than for the fluorescent reporters (Figures 3 and 5 and Figure S6). Note that while most anterior expression events are resolved in these data, a few of the ABa lineage *ref-2* expression domains are not (because the anterior vs posterior sister cells were not separately annotated at the stage when *ref-2* is expressed). However, the data that are resolved for *ref-2* are consistent with the promoter and fosmid reporter expression patterns. Gray branches on lineage trees indicate cells for which distinct transcriptomes could not be determined.

**Figure S6:**
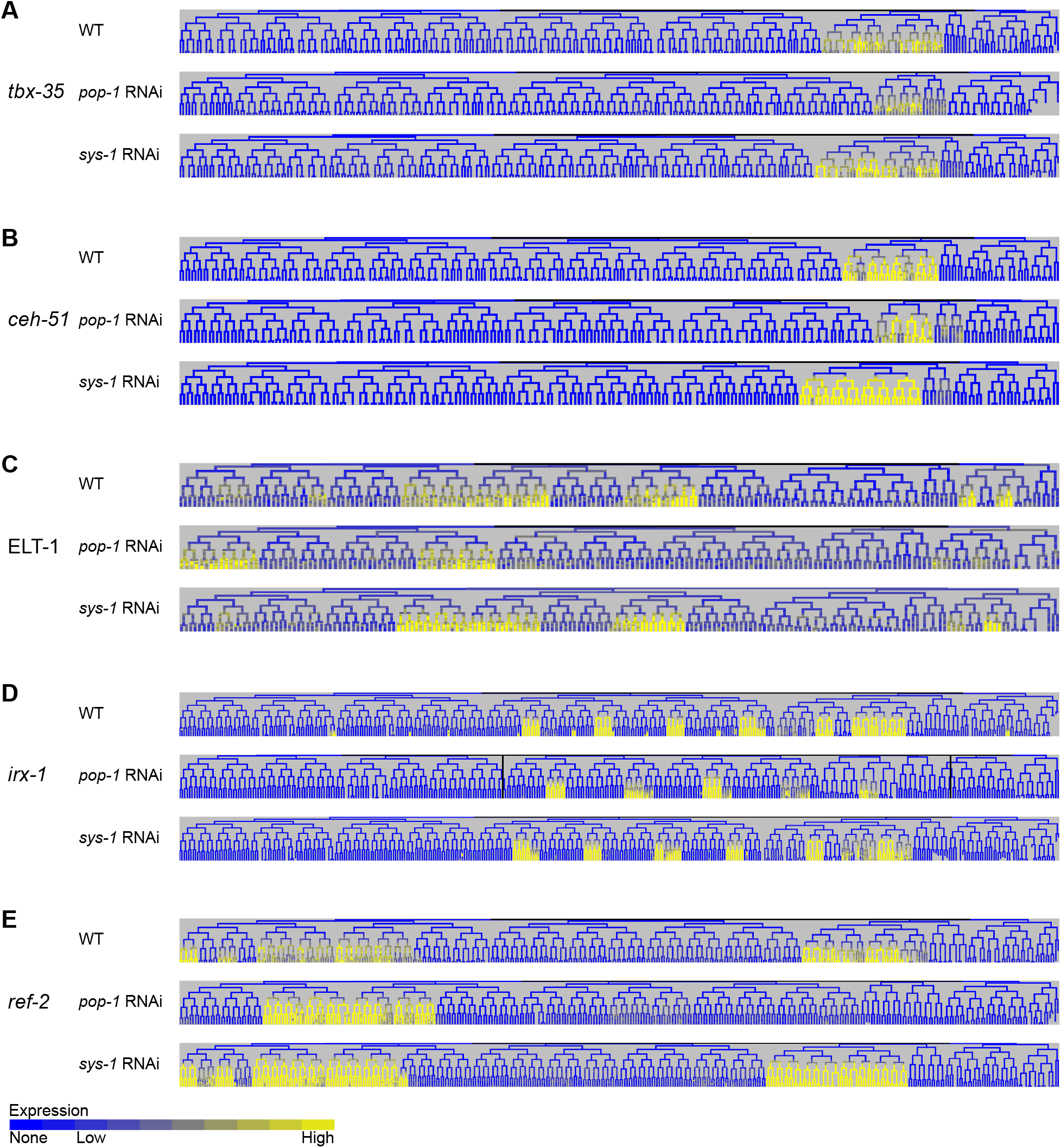
Anterior genes are regulated by Wnt effectors *pop-1* and *sys-1*. Full example WT, *pop-1* RNAi, and *sys-1* RNAi lineages are shown for integrated transgenic promoter reporters of *tbx-35* (A), *ceh-51* (B), *irx-1* (D), and *ref-2* (E) and for an integrated transgenic protein reporter of ELT-1 (C).

**Figure S7:**
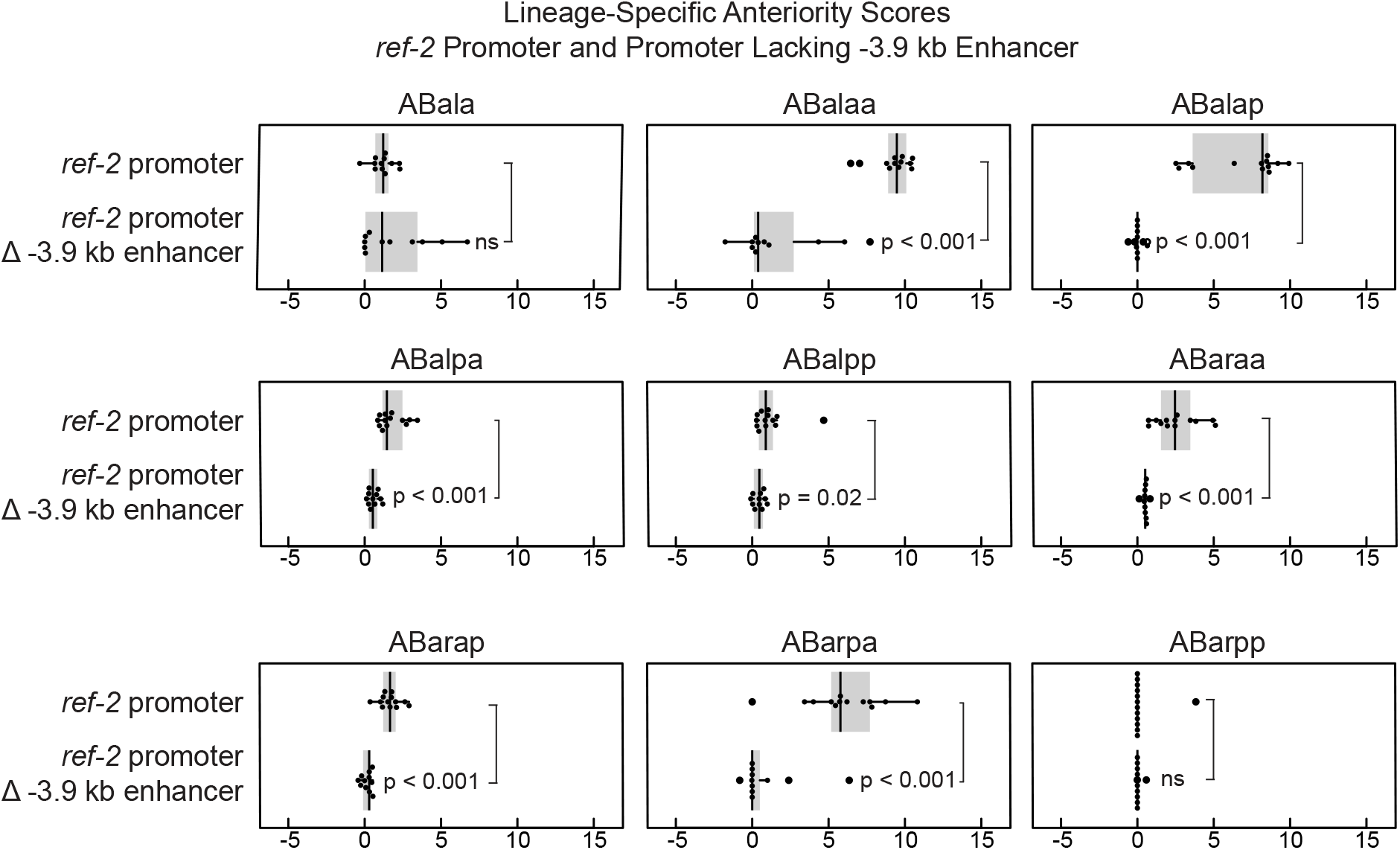
Deletion of the −3.9 kb enhancer from the *ref-2* promoter causes a reduction in the anterior bias in the expression driven by the promoter in most ABa sublineages in which the full-length promoter drives anterior-biased expression. Box plots display anteriority scores of lineages descended from indicated cells.

**Figure S8:**
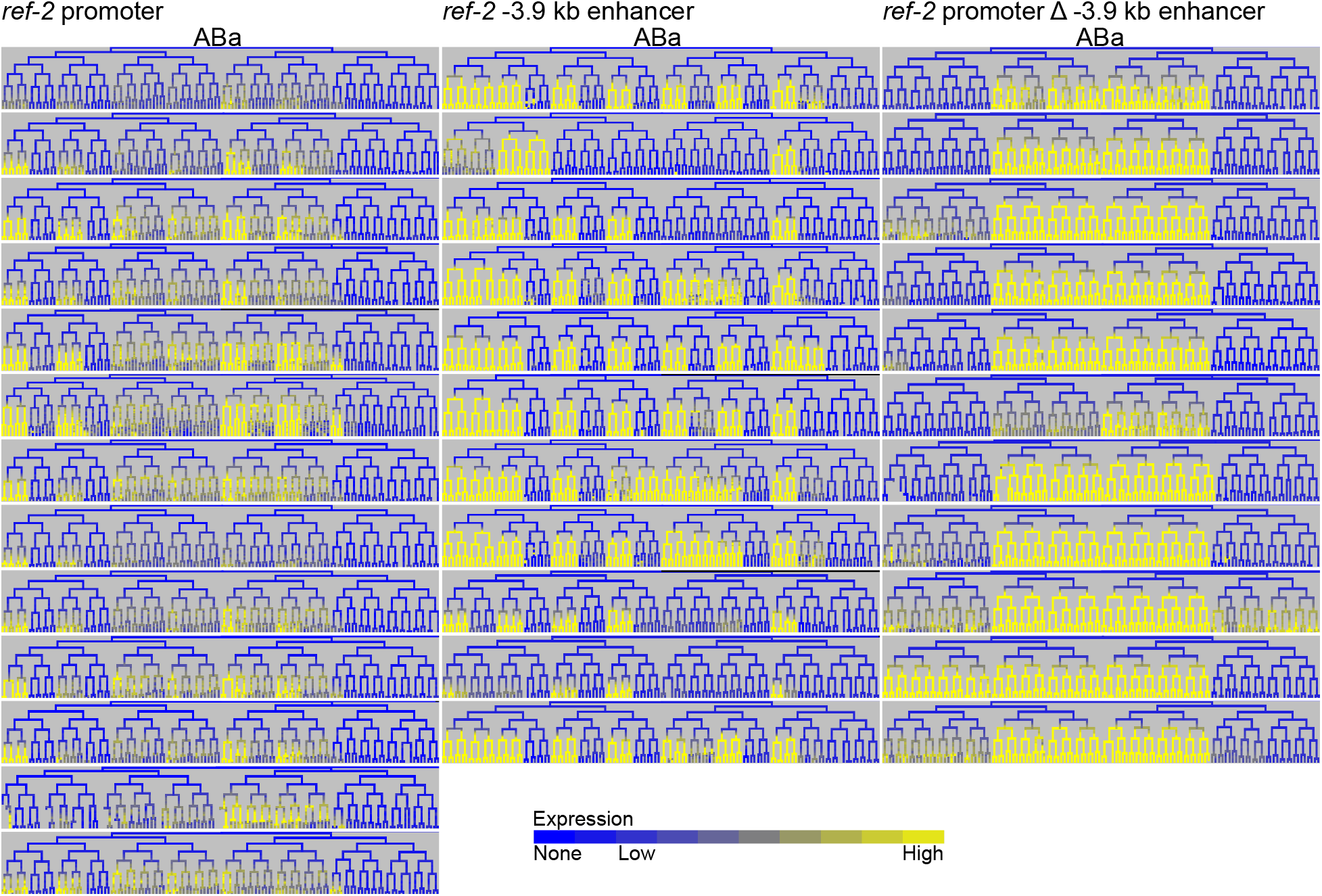
Expression patterns driven by the *ref-2* promoter, the −3.9 kb enhancer, and the promoter lacking the −3.9 kb enhancer are somewhat variable. Displayed here are partial lineages for all analyzed embryos bearing these reporters to show the variation in their expression.

**Figure S9:**
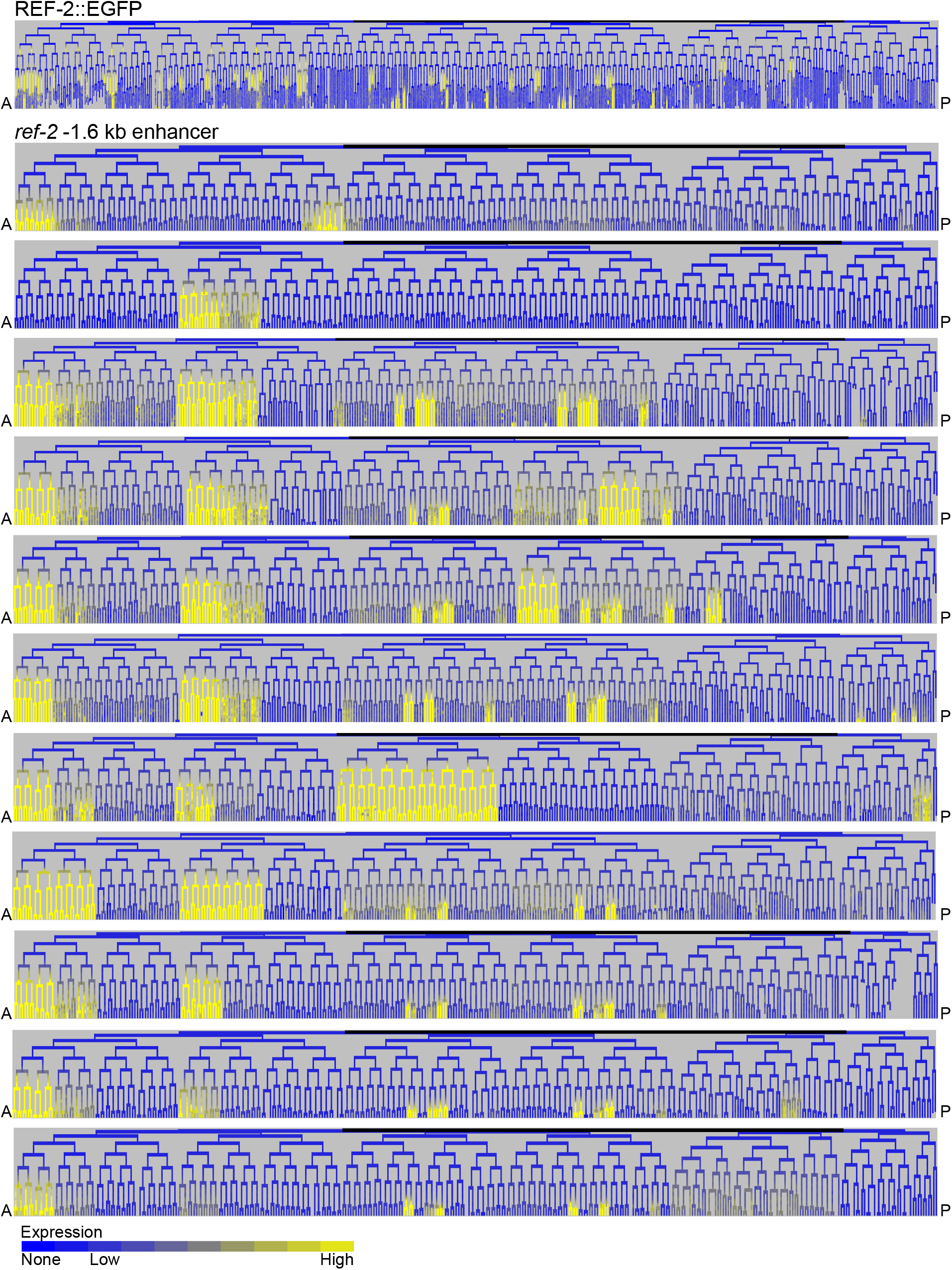
Expression patterns driven by the *ref-2* −1.6 kb enhancer are variable. Displayed here are full lineages for all analyzed embryos bearing this reporter to show the variation in its expression. The full lineage expression pattern of the REF-2::GFP protein reporter is shown for comparison.

**Figure S10:**
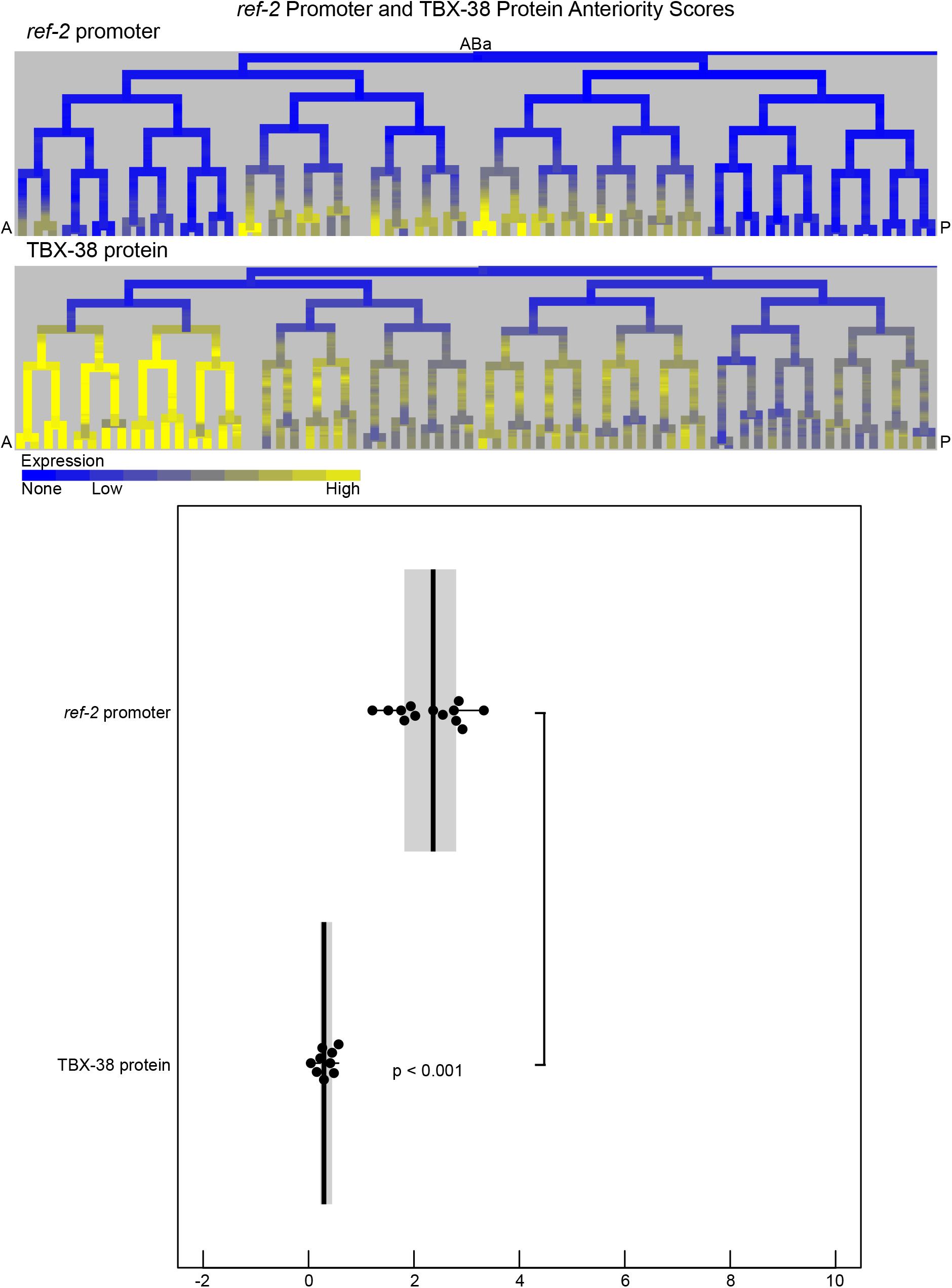
A TBX-38::GFP CRISPR knock-in protein reporter has minimally anterior-biased expression in the lineages in which the *ref-2* promoter drives anterior-biased expression. Displayed are partial lineages of the P*ref-2*::*his-24*::mCherry promoter reporter expression and the TBX-38::GFP CRISPR knock-in expression. The box plot shows the mean anteriority scores for the *ref-2* promoter reporter and the TBX-38::GFP CRISPR knock-in for the same lineages as in Figure 5J.

**Figure S11:**
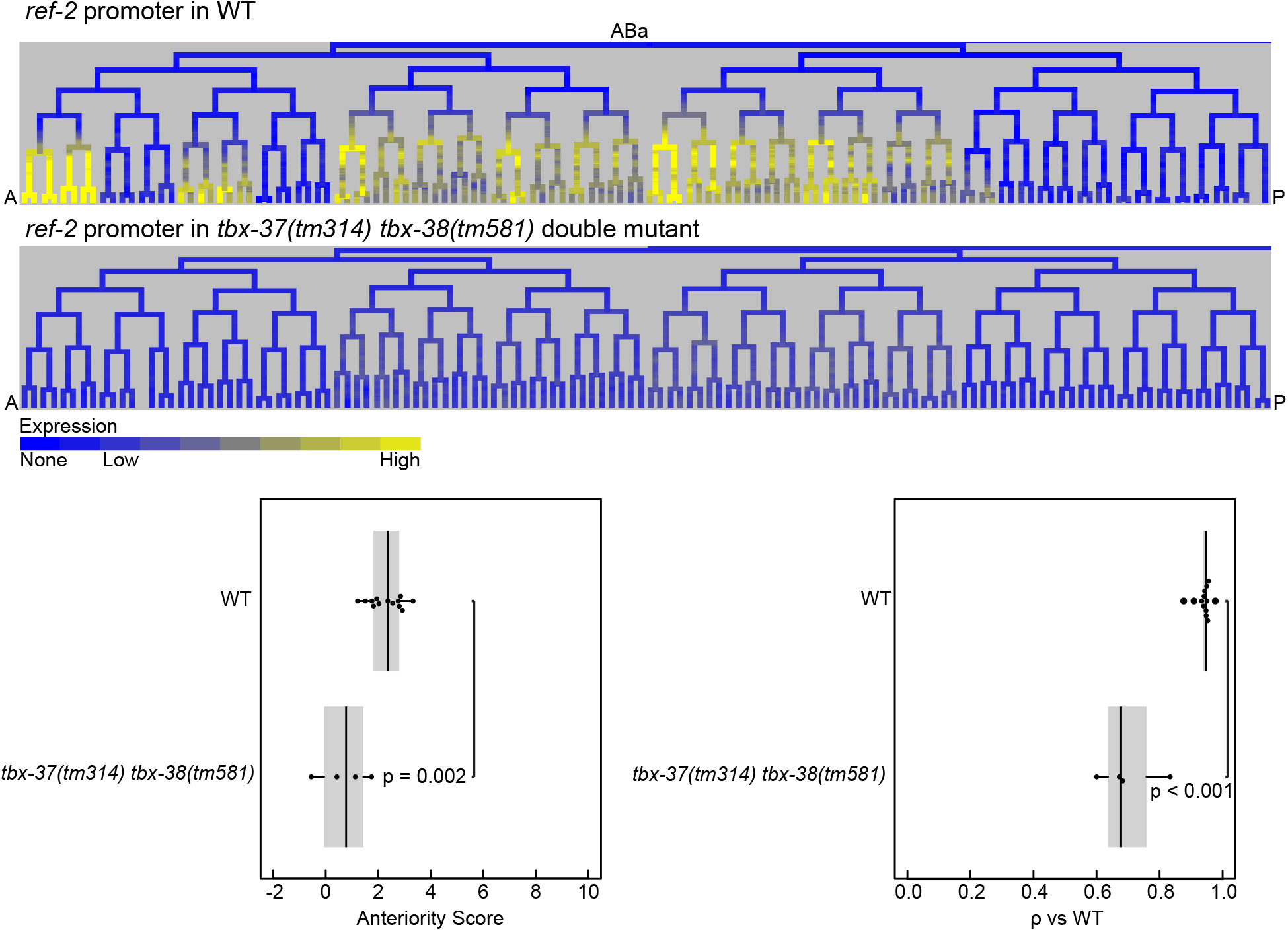
*tbx-37* and *tbx-38* are required for *ref-2* promoter-driven anterior-biased expression in the ABa lineage. Displayed are partial lineages for the expression driven by the *ref-2* promoter in a WT background and in a *tbx-37(tm314) tbx-38(tm581)* double mutant background. Also shown are box plots of anteriority scores and Spearman’s ρ for each of these groups, indicating a loss of anterior bias and correlation to WT for the *ref-2* promoter expression pattern in *tbx-37 tbx-38* double mutant embryos. Lineages used to determine anteriority scores are the same as in Figure 5J Spearman’s ρ analysis uses the full ABa lineage expression pattern and was calculated relative to the *ref-2* promoter in the WT background.

**Figure S12:**
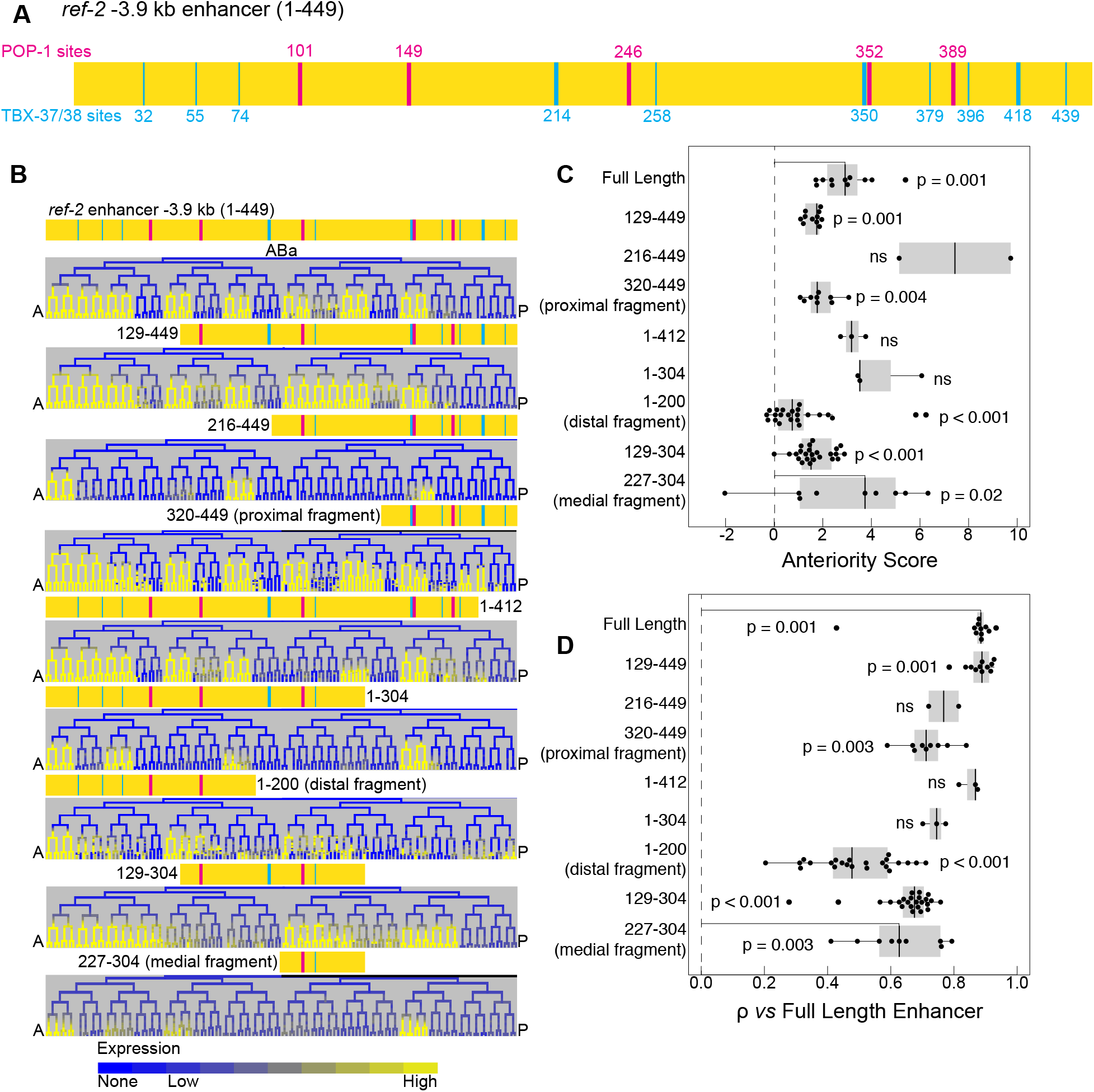
Three non-overlapping fragments of the −3.9 kb enhancer are each sufficient to drive anterior-biased expression in the ABa lineage of the early embryo. A) Model of the *ref-2* −3.9 kb enhancer with predicted TBX-37/38 and POP-1 sites indicated as in Figure 6. B) Expression patterns driven by full-length *ref-2* −3.9 kb enhancer and by all tested fragments. C-D) Box plots displaying the anteriority scores (C) and Spearman’s ρ (D) for the full-length *ref-2* −3.9 kb enhancer and all tested fragments. Lineages used to determine anteriority scores are the same as in Figure 5J. Spearman’s ρ analysis uses the full ABa lineage and was calculated relative to the full-length −3.9 kb enhancer.

**Figure S13:**
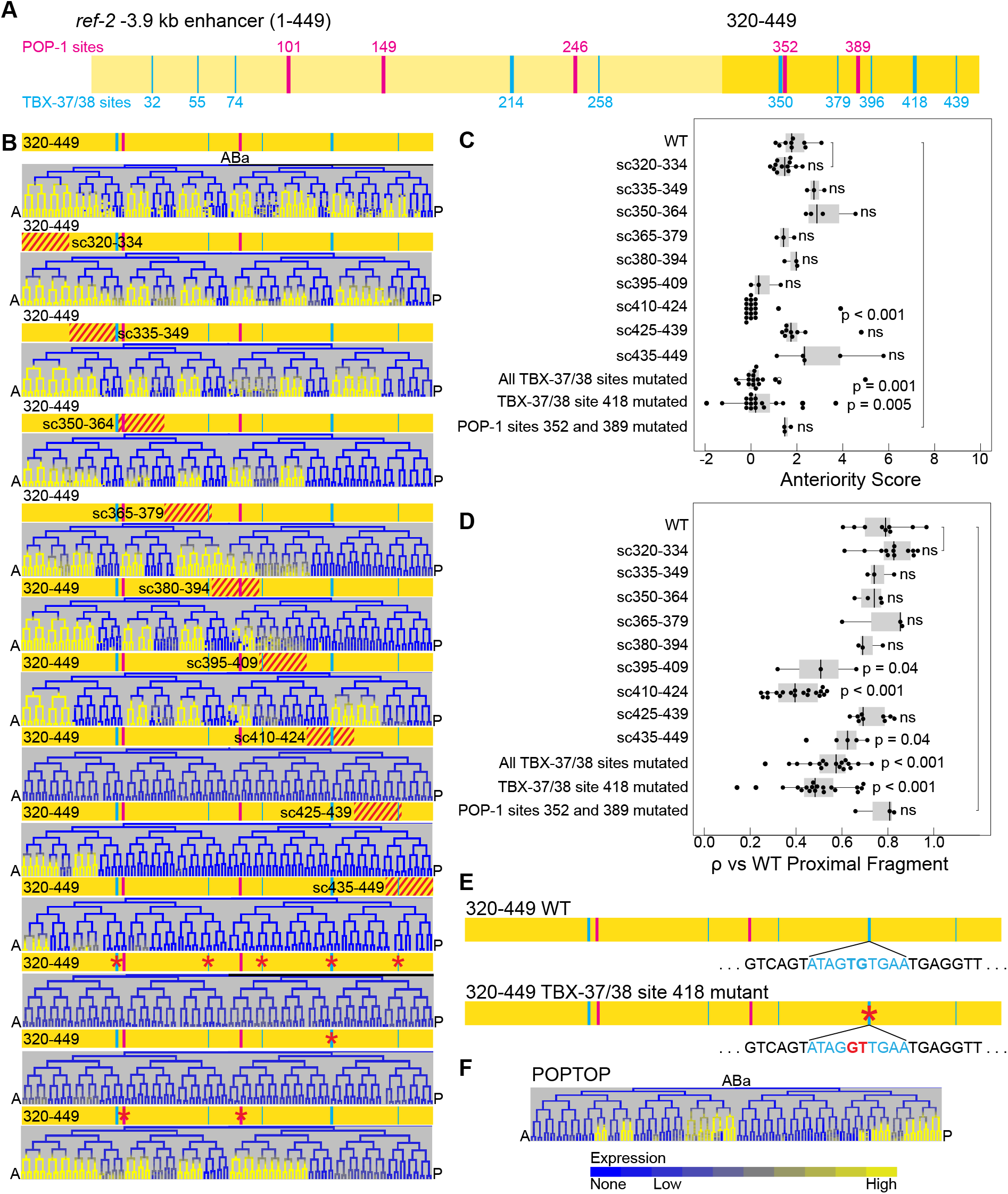
The broadly-expressed transcription factors *tbx-37* and *tbx-38* are required for the anterior-biased expression of *ref-2* in the ABa lineage. A) Model of the *ref-2* −3.9 kb enhancer with proximal fragment (base pairs 320-449) highlighted. Predicted TBX-37/38 and POP-1 sites are indicated as in Figure 6. B) Expression patterns driven by the proximal fragment of the *ref-2* −3.9 kb enhancer with its WT sequence, with each 15 base-pair segment of the fragment scrambled, with all predicted TBX-37/38 sites mutated, with *TBX_418_* mutated, and with the two predicted POP-1 sites mutated. C-D) Box plots displaying the anteriority scores (C) and Spearman’s ρ (D) for the proximal fragment with its WT sequence, with each 15 base-pair segment of the fragment scrambled, with all predicted TBX-37/38 sites mutated, with *TBX_418_* mutated, and with the two predicted POP-1 sites mutated. Lineages used to determine anteriority scores are the same as in Figure 5J. Spearman’s ρ analysis uses the full ABa lineage and was calculated relative to the WT proximal fragment. E) Models of the proximal fragment of the *ref-2* −3.9 kb enhancer with the WT sequence of *TBX_418_* and the mutated sequence used in the *TBX_418_* mutant reporters. F) Expression pattern driven by the seven-copy POP-1 site concatemer POPTOP [13,53].

**Figure S14:**
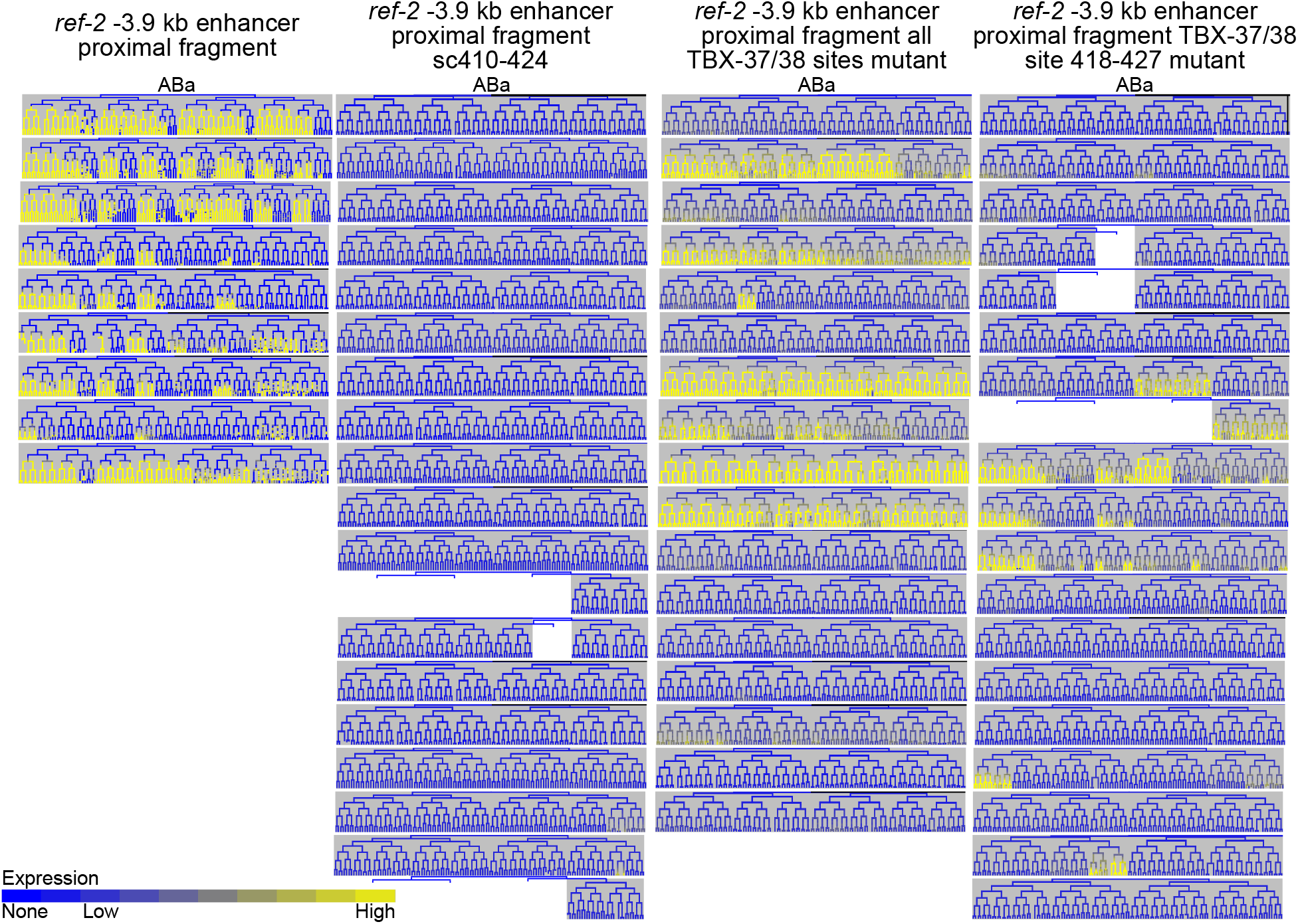
*ref-2* −3.9 kb enhancer WT proximal fragment, proximal fragment with all TBX-37/38 sites mutated, and proximal fragment with *TBX_418_* mutated exhibit variation in their expression patterns. Displayed are partial lineages for all analyzed embryos expressing reporters for the *ref-2* −3.9 kb enhancer WT proximal fragment, the *ref-2* −3.9 kb enhancer proximal fragment with base pairs 410-424 scrambled, the *ref-2* −3.9 kb enhancer proximal fragment with all TBX-37/38 sites mutated, and the *ref-2* −3.9 kb enhancer proximal fragment with *TBX_418_* mutated to show the variation in the expression pattern for each of these reporters.

## TABLES

Table S1:

*Caenorhabditis elegans* strain list

Table S2:

*Escherichia coli* strain list

Table S3:

Quantitative analyses raw data

Table S4:

Anterior gene summary expression data

Table S5:

Recombinant DNA list

## ACKNOWLEDGMENTS

We thank J. Archibald Millar, Julia Richards and Shaili Patel for technical contributions to data collection and analysis. We thank Meera Sundaram for use of her injection equipment. We thank Christopher Brown, Meera Sundaram, Michael Atchison, Peter Klein, and Mary Mullins for helpful discussions. Some strains were provided by the CGC, which is funded by NIH Office of Research Infrastructure Programs (P40 OD010440). We also thank the Waterston lab, the Maduro lab, and the Cochella lab for kindly providing *C. elegans* strains. We thank the *C. elegans* Reverse Genetics Core Facility at the University of British Columbia, part of the International *C. elegans* Gene Knockout Consortium, which produced the *ref-2* mutant allele (*ref-2(gk178))* used in this study. We thank the National Bioresource Project, Tokyo, Japan, which is part of the International *C. elegans* Gene Knockout Consortium, which produced the *tbx-37* and *tbx-38* mutant alleles (*tbx-37(tm314)* and *tbx-38(tm581)*) used in this study. This work was funded by T32GM008216, F31GM123737, and R35GM130357.

